# An auto-inhibitory mechanism regulates the non-enzymatic functions of a histone demethylase

**DOI:** 10.1101/545814

**Authors:** Gulzhan Raiymbek, Sojin An, Nidhi Khurana, Saarang Gopinath, Saikat Biswas, Ajay Larkin, Raymond Trievel, Uhn-soo Cho, Kaushik Ragunathan

## Abstract

H3K9 methylation (H3K9me) specifies the establishment and maintenance of transcriptionally silent epigenetic states or heterochromatin. The enzymatic erasure of histone modifications is widely assumed to be the primary mechanism that reverses epigenetic silencing. Here, we reveal an inversion of this paradigm where a putative histone demethylase Epe1 in fission yeast, has a non-enzymatic function that opposes heterochromatin assembly. An auto-inhibitory conformation regulates the non-enzymatic properties of Epe1 and licenses its interaction with Swi6^HP1^. Mutations that map to the putative catalytic JmjC domain preserve Epe1 in an auto-inhibited state which disrupts its localization to sites of heterochromatin formation and interaction with Swi6^HP1^. H3K9me relieves Epe1 from auto-inhibition and stimulates its ability to form an inhibitory complex with Swi6^HP1^ *in vitro* and in cells. Our work reveals that histone modifications, rather than being passive scaffolds, actively participate in regulating the assembly of specific epigenetic complexes in cells.

## INTRODUCTION

The covalent and reversible modification of histones allows cells to establish stable and heritable patterns of gene expression without any changes to their genetic blueprint. Histone H3 lysine 9 methylation (H3K9me) is a conserved mark of transcriptional silencing that is associated with the formation of specialized domains called heterochromatin. Heterochromatin formation is critical for centromere and telomere function, silencing of transposons and repetitive DNA elements and for maintaining lineage-specific patterns of gene expression (Grewal and Jia, 2007; Nicetto et al., 2019). It is widely assumed that a delicate interplay between readers, writers and erasers of histone marks regulate the establishment and maintenance of epigenetic states (Allis and Jenuwein, 2016). In fission yeast, the establishment of H3K9 methylation requires the enzymatic activity of a conserved methyltransferase, Clr4^Suv39h^ (Rea et al., 2000). H3K9 methylation subsequently recruits chromatin effector proteins with reader domains that recognize and bind specific histone marks. Two HP1 proteins, Swi6^HP1^ and Chp2^HP1^ are involved in this process and have distinct, non-overlapping functions during transcriptional silencing (Motamedi et al., 2008). Ultimately, factors such as Epe1, a putative histone demethylase, promotes epigenetic erasure thus reversing H3K9 methylation and transcriptional gene silencing {Trewick, 2005 #1409}.

The sequence-specific recruitment of Clr4^Suv39h^ restricts heterochromatin establishment to distinct sites in the genome such as the centromeres, telomeres and the mating type locus (Bayne et al., 2010; Hall et al., 2002; Verdel et al., 2004; Volpe et al., 2002). Once established, H3K9 methylation spreads to silence genes that are distant from heterochromatin nucleation centers. H3K9me spreading depends on the read-write activity of Clr4^Suv39h^ (Al-Sady et al., 2013; Zhang et al., 2008). Two conserved, structural attributes of Clr4^Suv39h^ mediate this process: a conserved chromodomain that recognizes and binds to H3K9me and an enzymatic SET domain that is involved in catalysis (Ivanova et al., 1998). The ability of Clr4^Suv39h^ to bind to the product of its own enzymatic activity enables H3K9 methylated histones to serve as carriers of epigenetic information (Audergon et al., 2015; Ragunathan et al., 2015). Following DNA replication, H3K9 methylated histones that are partitioned between daughter DNA strands serve as templates to mark newly deposited histones.

H3K9 methylation serves as a multivalent platform that recruits the HP1 homolog, Swi6^HP1^ to sites of constitutive heterochromatin (Ekwall et al., 1995; Ekwall et al., 1996). HP1 proteins have a conserved architecture consisting of a chromodomain (CD) that is involved in H3K9 methylation binding and a dimerization domain called the chromoshadow domain (CSD) that mediates protein-protein interactions (Bannister et al., 2001; Cowieson et al., 2000). The oligomerization of HP1 proteins promotes the formation of higher order chromatin complexes that exhibit phase separated, liquid-like properties *in vitro* and in cells (Larson et al., 2017; Sanulli et al., 2018; Strom et al., 2017). Swi6^HP1^ interacts with a broad spectrum of agonists and antagonists that influence epigenetic silencing. Most notably, the recruitment of Epe1, a putative histone demethylase that opposes heterochromatin formation, is dependent on Swi6^HP1^ (Ayoub et al., 2003). The loss of Epe1 leads to increased heterochromatin spreading beyond normal boundary sequences, the inheritance of H3K9 methylation via sequence-independent pathways, increased cell to cell variation in H3K9 methylation patterns and the acquisition of adaptive epigenetic traits (Audergon et al., 2015; Ayoub et al., 2003; Ragunathan et al., 2015; Wang et al., 2015; Zofall and Grewal, 2006; Zofall et al., 2012). Despite its critical role in heterochromatin regulation, how Epe1 exerts its anti-silencing activity in cells remains mysterious (Trewick et al., 2007).

Histone demethylases have prominent roles in regulating the reversibility of epigenetic states (Iwase et al., 2007; Whetstine et al., 2006). Changes in their expression leads to widespread chromatin reorganization which alters both the prognosis and treatment of diseases such as cancer (Liau et al., 2017). Epe1 most closely resembles JumonjiC (JmjC) domain containing proteins that use Fe (II) and α-ketoglutarate as co-factors to catalyze histone demethylation (Tsukada et al., 2006). Unlike active histone demethylases where an HxD/E….H motif is involved in Fe (II) coordination, Epe1 has a non-canonical HXE….Y motif (Tsukada et al., 2006). Although Epe1 shares conserved features with other histone demethylases at the amino acid level, there is no biochemical evidence to support the notion that Epe1 has any *in vitro* enzymatic activity. Epe1 purified from fission yeast cells, or a recombinant source (insect cells) exhibits no H3K9 demethylase activity *in vitro* (Tsukada et al., 2006; Zofall and Grewal, 2006). However, point mutations of amino acid residues involved in Fe (II) or α-ketoglutarate binding negatively affect Epe1 activity in cells leading to heterochromatin spreading beyond normal boundary sequences (Trewick et al., 2007). These conflicting lines of biochemical and genetic data have prompted several alternative explanations for how Epe1 might exert its anti-silencing function in cells. This includes the possibility that Epe1 acts as a protein hydroxylase that targets non-histone proteins such as Swi6^HP1^, regulates the activity of the multi-subunit H3K9 methyltransferase CLRC complex, or functions as an H3K9 demethylase when in complex with Swi6^HP1^ (Aygun et al., 2013; Iglesias et al., 2018; Trewick et al., 2007; Zofall and Grewal, 2006). Although these models represent attractive possibilities for how Epe1 regulates heterochromatin, there is no direct evidence to suggest that any of these proteins represent bonafide enzymatic targets. An alternative hypothesis is that Epe1 has a non-enzymatic function that regulates heterochromatin spreading and epigenetic inheritance. In support of this hypothesis, the overexpression of Epe1 Fe (II) and α-ketoglutarate binding mutants suppresses heterochromatin spreading defects that are observed in Epe1 null cells (Trewick et al., 2007; Zofall and Grewal, 2006). These observations highlight that co-factor binding mutants of Epe1 can function as multi-copy suppressors despite the presumptive loss of any trace of enzymatic activity.

We discovered that the putative catalytic JmjC domain of Epe1 is at least in part, dispensable for its anti-silencing function in cells. The non-enzymatic function of Epe1 is regulated by auto-inhibition where the C-terminus of the protein interacts with its N-terminal half both in *cis* and in *trans.* Auto-inhibition is relieved in the presence of H3K9 methylation binding which stimulates complex formation between Epe1 and Swi6^HP1^. *In vivo*, this dependency on H3K9 methylation restricts the interaction between Epe1 and Swi6^HP1^ to sites of heterochromatin formation. This mode of selective complex formation enables sub-stoichiometric levels of Epe1 to effectively inhibit Swi6 mediated heterochromatin assembly through a non-enzymatic process. We propose that an auto-inhibitory mechanism regulates the non-enzymatic functions of Epe1 and highlights the versatile ways in which histone demethylases can oppose epigenetic inheritance.

## RESULTS

### A point mutation within the catalytic JmjC domain of Epe1 affects its localization at sites of constitutive heterochromatin

JmjC domain containing proteins require Fe (II) and α-ketoglutarate as co-factors to catalyze histone demethylation. Aligning the primary amino acid sequences of active histone demethylases with Epe1 reveals a naturally occurring histidine to tyrosine substitution (Y370) within a conserved triad of amino acid residues that coordinates iron (**Supplementary Figure 1A**). We tested whether this non-conserved tyrosine (Y370) is required for Epe1 activity in cells. To measure Epe1 activity, we used a reporter gene assay where the H3K9 methyltransferase, Clr4^Suv39h^ is fused to a DNA binding protein, TetR. This fusion protein is recruited to an ectopic site where ten Tet operator sites (*10X TetO*) are placed upstream of a reporter gene, *ade6^+^* (**Figure 1A**). Establishment in the absence of tetracycline results in the appearance of red colonies. The sequence-specific initiator, TetR-Clr4-I, dissociates in the presence of tetracycline enabling us to test whether cells can maintain silencing in the absence of continuous initiation. Wild-type cells are initially red on -tetracycline medium indicating that the reporter gene starts out being repressed (establishment). Wild-type cells turn white and exhibit no maintenance of epigenetic silencing when plated on +tetracycline containing medium. The ability of fission yeast cells to autonomously propagate epigenetic silencing across multiple generations is exquisitely sensitive to Epe1 activity. Epigenetic maintenance is observed in cells where Epe1 is either deleted or inactivated resulting in the appearance of red or sectored colonies on +tetracycline containing medium (Audergon et al., 2015; Ragunathan et al., 2015).

**Figure 1:**
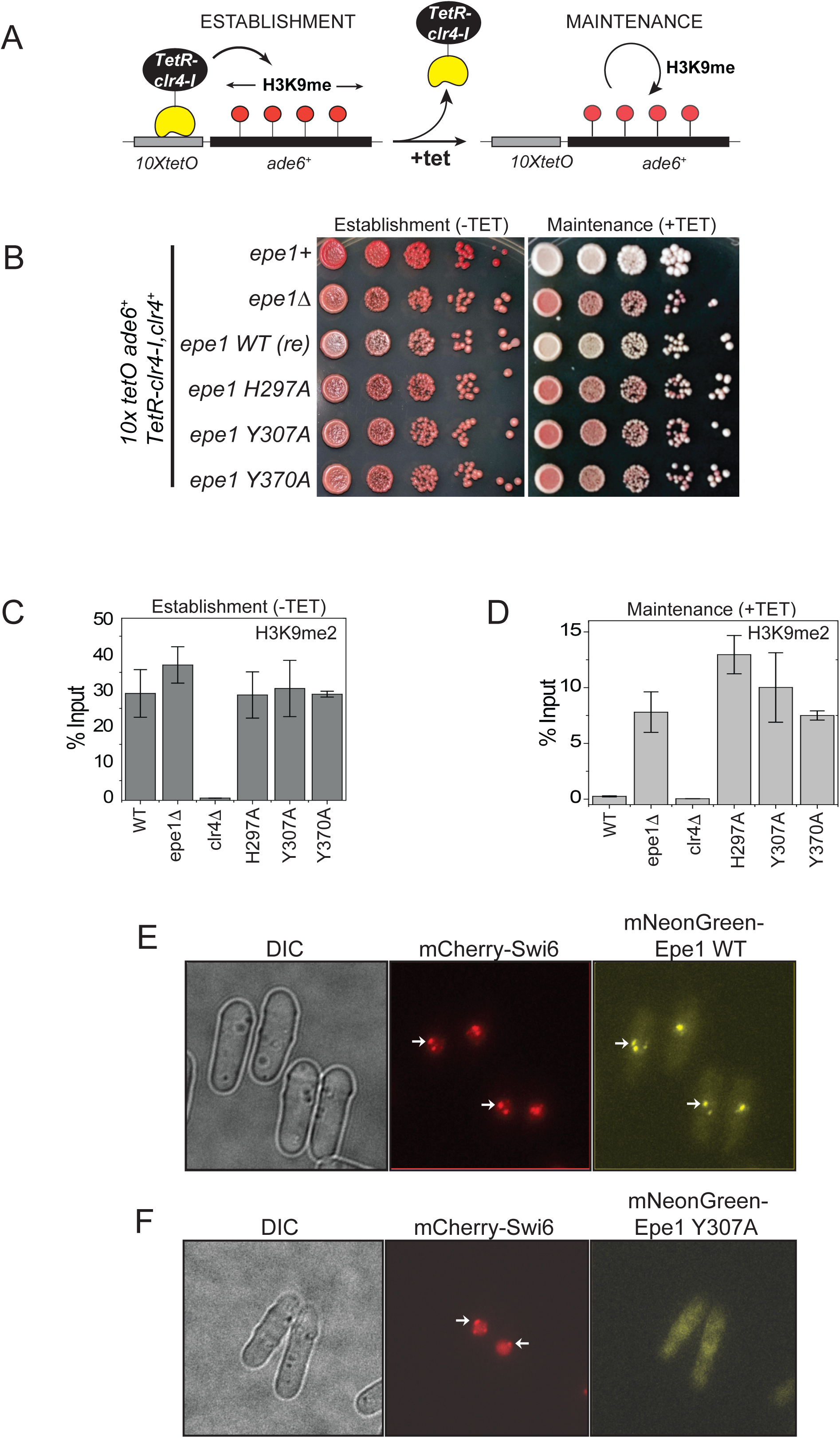
A point mutation within the catalytic JmjC domain of Epe1 affects protein localization at sites of constitutive heterochromatin. **A)** A reporter system to measure epigenetic inheritance. TetR-Clr4-I binding (-tetracycline) leads to the ectopic establishment of H3K9 methylation. The addition of tetracycline promotes TetR-Clr4-I dissociation allowing us to measure the extent of initiator independent maintenance of H3K9 methylation. **B)** A color based assay to detect the establishment and maintenance of epigenetic states. The establishment of epigenetic silencing (-tetracycline) leads to the appearance of red colonies. Epigenetic inheritance, indicated by red or sectored colonies, (+tetracycline) is critically dependent on Epe1 activity. Point mutations within the JmjC domain of Epe1 disrupt Epe1 activity leading to the appearance of red or sectored colonies. **C)** ChIP-qPCR measurements of H3K9me2 levels at the ectopic site (*10X tetO ade6+*) during establishment (-tetracycline) in different Epe1 mutant backgrounds (N=2). Error bars represent standard deviations. **D)** ChIP-qPCR measurements of H3K9me2 levels at the ectopic site (*10X tetO ade6+*) during maintenance (+tetracycline) in different Epe1 mutant backgrounds (N=2). Error bars represent standard deviations. **E)** Live cell imaging of Epe1 and Swi6. Three images in each case correspond to DIC, 488 excitation and 560 excitation channels. mNeonGreen-Epe1 and mCherry-Swi6^HP1^ form co-localized foci in green and red emission channels respectively (see white arrows). **F)** mNeonGreen-Epe1 Y307A fails to form any foci and instead exhibits a diffuse signal that permeates the nucleus. mCherry-Swi6^HP1^ forms foci corresponding to sites of constitutive heterochromatin.

Alanine substitutions of amino acid residues involved in Fe (II) or α-ketoglutarate binding (*epe1 H297A and epe1 Y307A* respectively) disrupt co-factor binding resulting in the concomitant loss of Epe1 activity. When expressed at endogenous levels, these mutant alleles form red or sectored colonies on +tetracycline containing medium and resemble *epe1Δ* cells (**Figure 1B**). Replacing the non-conserved tyrosine residue in Epe1 with alanine (*epe1 Y370A)* leads to a similar loss of function phenotype. Hence, despite the lack of conservation, a naturally tyrosine substitution within the JmjC domain of Epe1 is essential for its cellular activity. We used chromatin immunoprecipitation assays followed by qPCR to measure H3K9me2 levels associated with the reporter gene locus before and after tetracycline addition. Both wild-type and Epe1 mutant strains exhibit high levels of H3K9me2 during establishment (**Figure 1C**). However, wild-type cells lose H3K9me2 approximately twenty four hours after tetracycline addition. In contrast, Epe1 mutants that exhibit a red or sectored phenotype upon +tetracycline addition retain high levels of H3K9 methylation at the ectopic site (**Figure 1D**). We verified that the expression level of all Epe1 mutant proteins is equal relative to an actin loading control. Hence, neither overexpression artifacts nor changes in protein stability contribute to the maintenance specific phenotype we observed in our genetic assays (**Supplementary Figure 1B**).

Epe1 is stabilized at sites of constitutive heterochromatin through its interactions with Swi6^HP1^ (Ayoub et al., 2003; Zofall and Grewal, 2006). We imaged Epe1 and Swi6^HP1^ in live fission yeast cells using fluorescent protein fusions. We labeled Epe1 with mNeonGreen and Swi6^HP1^ with mCherry. This labeling scheme allows Epe1 and Swi6^HP1^ to be visualized in separate green and red emission channels respectively. Both fusion proteins were expressed from their endogenous promoters to discount any possible overexpression artifacts. mCherry-Swi6^HP1^ typically exhibits two or three bright foci in individual cells corresponding to sites of constitutive heterochromatin (centromeres and telomeres). mNeonGreen-Epe1 co-localizes with mCherry-Swi6^HP1^ as evidenced by the significant overlap between the bright foci that appear in both the green and red emission channels (see white arrows) (**Figure 1E**). Surprisingly, an Epe1 co-factor binding mutant, mNeonGreen-Epe1 Y307A, fails to co-localize with mCherry-Swi6^HP1^. Instead, the mutant protein exhibits a diffuse green signal within the nucleus and a complete lack of nuclear foci that co-localize with Swi6^HP1^ (**Figure 1F**). Hence, in addition to affecting any putative enzymatic functions, a co-factor binding mutation within the JmjC domain of Epe1 completely eliminates its localization at sites of constitutive heterochromatin.

### Mutations within the JmjC domain disrupt a direct interaction between Epe1 and Swi6^HP1^

We hypothesized that the absence of heterochromatin localization in the Epe1 JmjC mutant could reflect a loss of Swi6^HP1^ binding. We used a co-immunoprecipitation assay to compare the interaction between Epe1 and Swi6^HP1^ in wild-type and Epe1 mutant cells. We expressed an Epe1-3X FLAG fusion protein at endogenous levels. Using a FLAG antibody, we pulled-down Epe1 and detected its interaction with Swi6^HP1^ using a Swi6^HP1^ specific primary antibody. Swi6^HP1^ is enriched in pull-down experiments in wild-type cells relative to an untagged control (**Figure 2A)**. However, mutations in residues that affect Fe(II) or α-ketoglutarate binding (H297A, Y307A and Y370A) significantly attenuate this interaction (**Figure 2A)**. Interestingly, the Epe1 H297A mutant exhibits some residual interaction which could explain why its overexpression might suppress phenotypes observed in *epe1Δ* cells despite the loss of enzymatic activity. Our co-immunoprecipitation assays reveal that alanine substitutions of nearly every residue that is implicated in co-factor binding, eliminates the binding between Epe1 and Swi6^HP1^.

**Figure 2:**
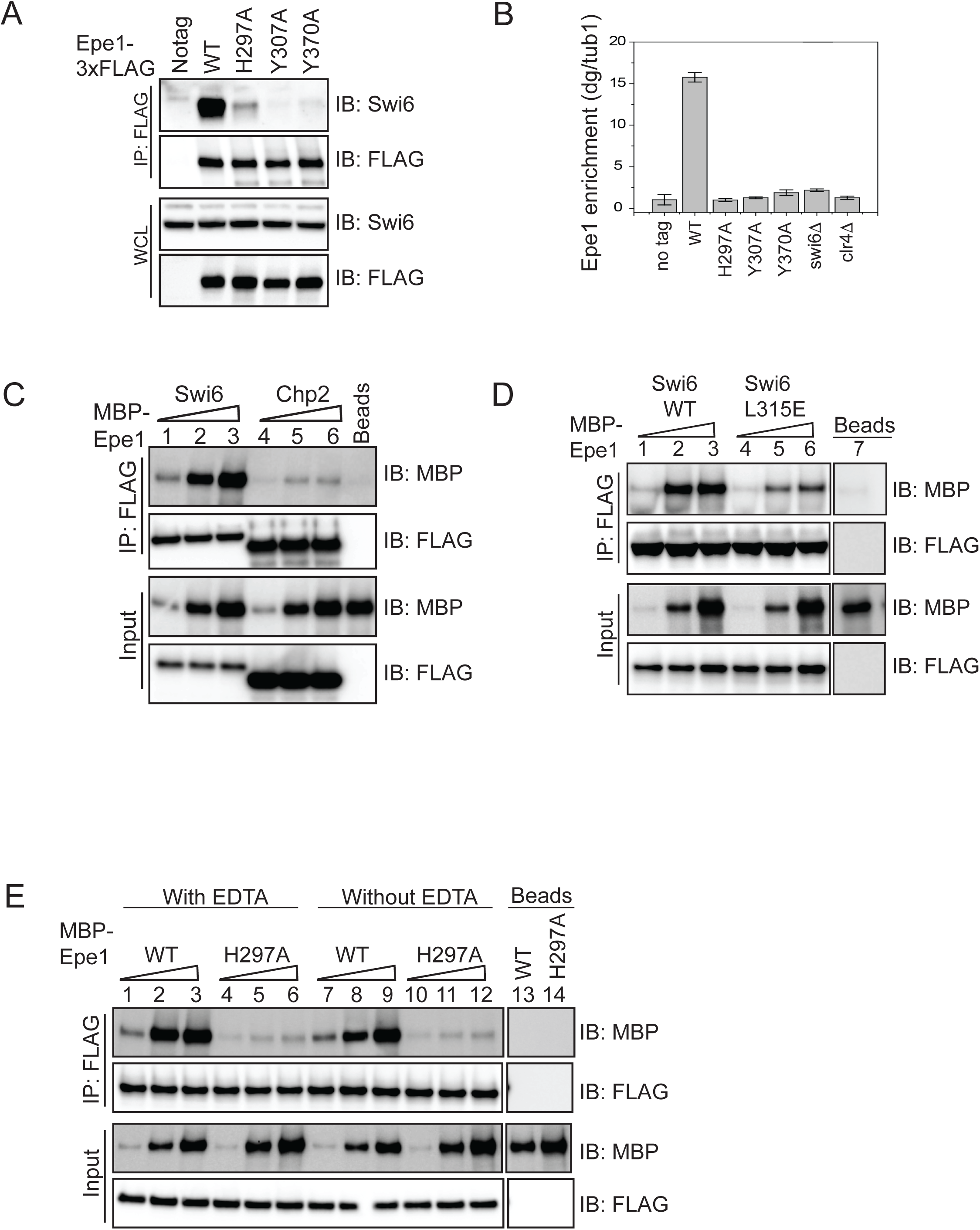
Mutations within the JmjC domain perturb a direct interaction between Epe1 and Swi6^HP1^. **A)** Western blots of co-immunoprecipitation measurements to test the interaction between Epe1-3XFLAG and Swi6^HP1^. Epe1 is detected using a FLAG antibody and Swi6^HP1^ is detected using a primary antibody. The interaction between the two proteins is preserved in wild-type cells and is completely eliminated in most Epe1 JmjC mutants. **B)** ChIP-qPCR measurements of Epe1 occupancy at sites of constitutive heterochromatin (*dg* pericentromeric repeats) (N=2). Error bars represent standard deviations. Epe1 enrichment is reduced to near background levels (*swi6Δ* and *clr4Δ*) in Epe1 mutants that exhibit a loss of activity. **C)** Western blots of *in vitro* binding assays using recombinant Epe1 protein. Increasing amounts of wild-type MBP-Epe1 protein are added while maintaining a fixed amount of 3X-FLAG-Swi6^HP1^ or 3X-FLAG-Chp2 ^HP1^ on beads. Epe1 exhibits a strong preference for Swi6^HP1^over Chp2 ^HP1^. **D)** Increasing amounts of wild-type MBP-Epe1 protein are added while maintaining a fixed amount of 3X-FLAG-Swi6^HP1^ or a chromoshadow domain (CSD) mutant, 3X-FLAG Swi6^HP1^ L315E on beads. Western blots reveal that a point mutation in the conserved Swi6 ^HP1^ chromoshadow domain (L315E) leads to reduced levels of interaction between recombinant Epe1 and Swi6^HP1^ L315E. **E)** Increasing amounts of wild-type MBP-Epe1 and MBP-Epe1 H297A are added while maintaining a fixed amount of 3XFLAG-Swi6^HP1^ on beads. Experiments were performed in the presence and absence EDTA to measure co-factor independent interactions between the two proteins. Epe1 H297A binding to Swi6^HP1^ is significantly attenuated relative to the wild-type protein.

Epe1 is specifically enriched at sites of constitutive heterochromatin such as the pericentromeric *dg* and *dh* repeats, the mating type locus and the telomeres. We used formaldehyde to crosslink cells followed by chromatin immunoprecipitation to compare heterochromatin occupancy differences between Epe1 wild-type and Epe1 co-factor binding mutants. After crosslinking, we used a FLAG antibody to pull down the chromatin bound fraction of Epe1. We used qPCR to measure Epe1 occupancy at the pericentromeric *dg* repeats which are sites of constitutive heterochromatin formation in fission yeast. Epe1 is enriched within *dg* repeats in wild-type cells and this heterochromatin specific occupancy pattern is disrupted both in *swi6Δ* and *clr4Δ* cells (**Figure 2B)**. Hence, the localization of Epe1 to sites of heterochromatin formation depends on Swi6^HP1^ and H3K9 methylation. Mutations within the JmjC domain that disrupt co-factor binding lead to a substantial reduction in Epe1 occupancy at sites of heterochromatin formation. Consistent with our co-immunoprecipitation studies, all co-factor binding mutants of Epe1 exhibit a significant reduction or completely fail to localize to the pericentromeric *dg* repeats (**Figure 2B)**. The chromatin occupancy of Epe1 mutants as measured by ChIP-qPCR is strikingly similar to that of wild-type Epe1 in a heterochromatin deficient background (*swi6Δ* or *clr4Δ*). We altered our fixation conditions using additional reactive cross-linkers and extended the time for formaldehyde cross-linking. Altering cross-linking conditions did not result in any increase in our measurements of chromatin occupancy amongst Epe1 mutants (**Supplementary Figure 2A)**. Based on these results, we concluded that Epe1 co-factor binding mutants exhibit significant defects in their ability to interact with Swi6^HP1^ and a complete inability to localize to sites of heterochromatin formation.

Our co-immunoprecipitation, imaging and ChIP experiments preclude us from making any conclusions as to whether the interaction between Epe1 and Swi6^HP1^ is mutation dependent or requires the putative catalytic functions of Epe1 *in vivo*. To address this concern, we purified MBP fusions of wild-type Epe1 and Epe1 H297A from insect cells. Swi6^HP1^ was purified from *E.coli*. We used TEV protease to cleave the MBP tag and confirmed that Epe1 remains soluble but preserved the tag in subsequent purifications for our binding assays (**Supplementary Figure 2B**). We compared the thermal stability of the wild-type and mutant Epe1 protein (Epe1 H297A) using isothermal calorimetry measurements (**Supplementary Figure 2C**). Wild-type Epe1 and Epe1 H297A exhibit similar denaturation temperatures implying that the mutation within the JmjC domain does not destabilize the protein or lead to gross alterations in protein structure. The difference in peak intensities in the ITC profile merely reflects slightly different protein amounts used in this assay. These results are consistent with structural studies of JmjC domain containing proteins where the presence of co-factors within the active site does not substantially alter protein conformation {Horton, 2011 #1525}.

To perform *in vitro* binding assays, we immobilized Swi6^HP1^ on FLAG beads and added three different concentrations of Epe1. Epe1 was detected in these binding assays using an MBP antibody and the total amount of Swi6^HP1^ was measured using a FLAG antibody. Through a series of titration measurements we found that using an MBP antibody and a chemiluminesence based readout produces a limited linear response which precludes us from reporting an apparent Kd (data not shown). We verified that the Epe1 protein we purified from insect cells interacts with Swi6^HP1^ but not with a second HP1 homolog in *S.pombe*, Chp2^HP1^ (**Figure 2C**). Hence, the recombinant Epe1 protein we purified from insect cells recapitulates the known binding preference of Epe1 towards Swi6^HP1^ (Sadaie et al., 2008). The chromoshadow domain (CSD) of Swi6^HP1^ mediates protein dimerization and regulates Swi6^HP1^ dependent protein-protein interactions (Canzio et al., 2013). We expressed and purified a dimerization deficient mutant of Swi6 from *E.coli* (3XFLAG-Swi6^HP1^ L315E). Our binding assays using the mutant Swi6^HP1^ protein (Swi6^HP1^ L315E) reveals a significant reduction in its ability to interact with Epe1^WT^ (**Figure 2D**). Therefore, Epe1 interacts with Swi6^HP1^ through a conserved mechanism that is simultaneously shared across different heterochromatin associated binding proteins.

To test whether a mutation within the JmjC domain leads to a loss of interaction between Epe1 and Swi6^HP1^, we compared binding assays between the recombinant wild-type Epe1 protein and the Fe (II) binding deficient mutant, Epe1 H297A. Adding increasing amounts of the wild-type MBP-Epe1 leads to a corresponding increase in the amount of protein that interacts with Swi6^HP1^. However, this type of interaction and increase in binding is not observed in the case of MBP-Epe1 H297A (**Figure 2E**). We compared binding assays performed in the presence and absence of EDTA to rule out any potential contributions that may arise from divalent metals ions that interact with the JmjC domain. The interaction between Epe1 and Swi6^HP1^ is nearly identical in the presence or absence of EDTA (**Figure 2E)**. These results suggest that the interaction between Epe1 and Swi6^HP1^ is direct but dependent on mutations that map to the putative catalytic JmjC domain.

To test whether the enzymatic activity of Epe1 may enhance its interaction with Swi6^HP1^, we added both Fe (II), α−ketoglutarate and ascorbate to mimic “histone-demethylase reaction conditions” in our *in vitro* binding assays (Tsukada and Nakayama, 2010). The addition of co-factors required for histone demethylation did not dramatically alter the extent of interaction properties between Epe1 and Swi6^HP1^ (**Supplementary Figure 2D**). Hence, our *in vitro* assays fail to capture any effect that co-factor binding itself may have on the interaction between Epe1 and Swi6^HP1^. We also tested whether the interaction between *S.pombe* Swi6^HP1^ and Epe1 might be different from those we obtained using Swi6^HP1^ purified from *E.coli* (**Supplementary Figure 2E**). However, we observed no differences in the interaction profile suggesting that the loss of binding we detected in the Epe1 H297A mutant is independent of post-translational modifications (such as phosphorylation) that may alter the biochemical properties of Swi6^HP1^ (Shimada et al., 2009). These results suggest that mutations within the JmjC domain of Epe1 likely induce a dominant conformational change that negatively impacts Swi6^HP1^ binding.

We tested whether Swi6^HP1^ binding may activate the latent enzymatic properties of Epe1. We performed histone demethylase assays using recombinant Epe1 in the presence and absence of Swi6^HP1^. We used a histone H3 tail peptide with a tri-methyl modification at the lysine 9 position (H3K9me3 peptide) as a substrate. We were unable to detect a mass shift corresponding to the removal of one or more methyl groups in reactions that we performed with Epe1 alone or Epe1 in complex with a five-fold molar excess of Swi6 (**Supplementary Figure 2F**). In contrast, JMJD2A, an active demethylase, is fully capable of demethylating an H3K9me3 peptide substrate (**Supplementary Figure 2G**). Hence, Swi6^HP1^ binding to Epe1 is not sufficient to activate its putative enzymatic activity.

### Swi6^HP1^ interacts with the C-terminus of Epe1 through a region that is proximal to the JmjC domain

To map the Swi6^HP1^ interaction site within Epe1, we used an *in vitro* translation (IVT) assay where we expressed fragments of Epe1 and tested their ability to interact with Swi6^HP1^. We used a computational disorder prediction program to define ordered and disordered regions within Epe1 (**Supplementary Figure 3A**). The JmjC domain emerges as one of two ordered regions extending from amino acids 233-434. The second ordered domain that is located within the C-terminus of the protein has no known similarity to existing proteins structures, does not have any ascribed function but is conserved within the *Schizosaccharomyces* lineage (data not shown). We designed and expressed partial fragments of Epe1 using rabbit reticulocyte lysates including the full length protein as a positive control. We added FLAG beads that were pre-incubated with 3XFLAG-Swi6^HP1^ to the IVT extract. A C-terminal fragment of Epe1 spanning 434-948 amino acids and an N-terminal fragment of Epe1 encompassing 1-600 amino acids emerge as the strongest Swi6^HP1^ interaction candidates (**Supplementary Figure 3B)**. Based on this analysis, we concluded that the minimal Swi6^HP1^ binding site within Epe1 maps to a region between 434-600 amino acids. This region is proximal to but non-overlapping with the predicted JmjC domain of Epe1.

To validate the conclusions of our IVT binding assay, we expressed and purified both C-terminal fragments of Epe1 from *E.coli.* The first fragment encompasses the entire C-terminus of Epe1 from 434-948 amino acids (Epe1^434-948^). The second fragment corresponds to only the minimal Swi6^HP1^ interaction site spanning 434-600 amino acids (Epe1^434-600^). We performed binding assays comparing the relative binding strength of the full-length Epe1 protein and the two Epe1 fragments-Epe1^434-600^ and Epe1^434-948^ with Swi6^HP1^. Both Epe1 fragments exhibit an increase in their ability to interact with Swi6^HP1^ relative to the full-length protein (**Figure 3A-B**). Hence, the same amino acid sequences, when placed in the context of full length Epe1, are less accessible to Swi6^HP1^. These results raise the possibility that the JmjC domain has a steric function and its presence within the full length protein may inhibit Swi6^HP1^ binding. We also performed co-immunoprecipitation experiments in cells expressing 3XFLAG-Epe1^434-948^ and detected the same type of interaction with Swi6^HP1^ that we detected in our *in vitro* assay (**Supplementary Figure 3C**). We note that the expression level of the 3X FLAG-Epe1^434-948^ protein is substantially lower compared to full-length Epe1 protein with at least a four-fold lower expression level (**Supplementary Figure 3D**).

**Figure 3:**
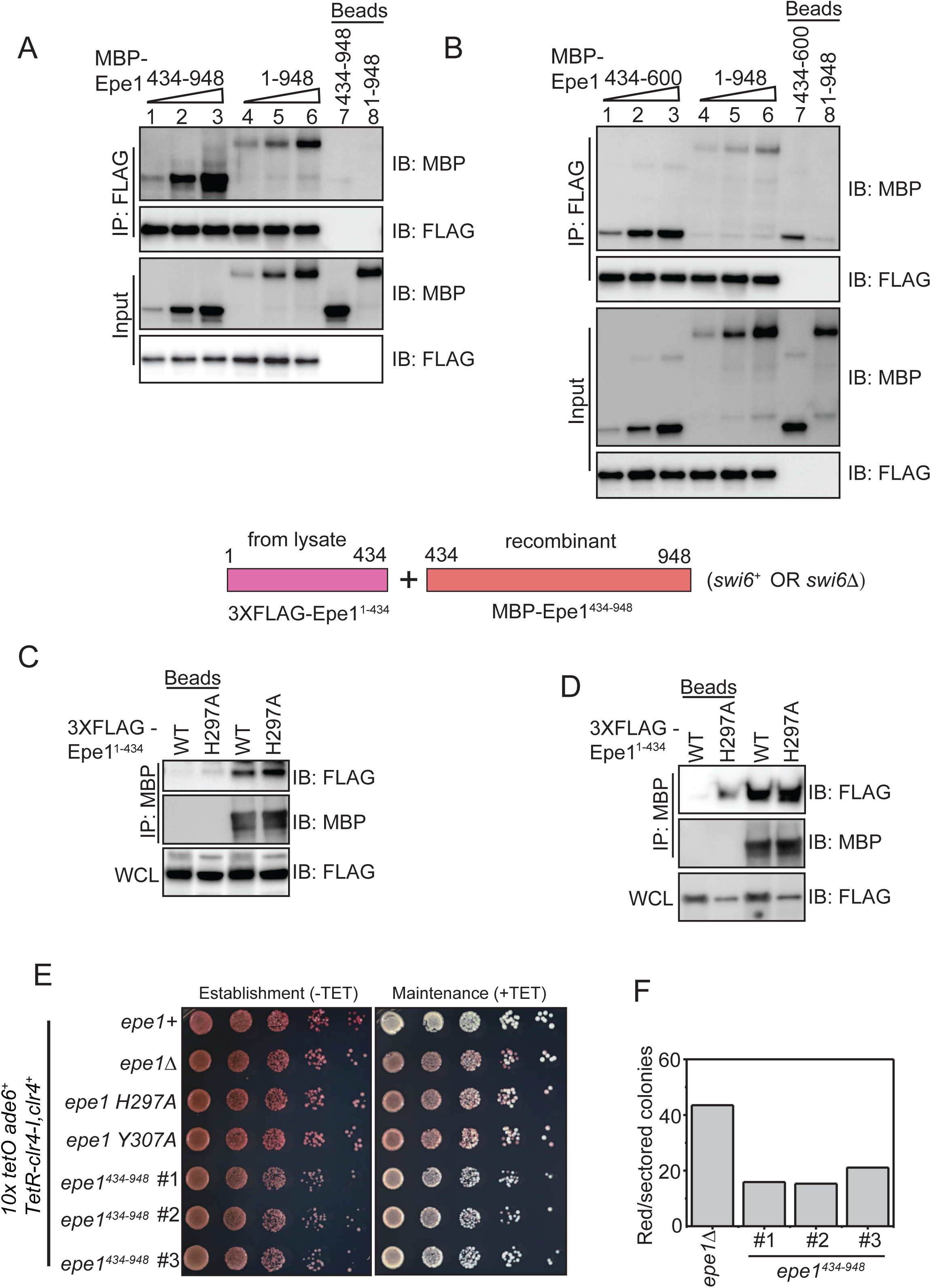
Swi6^HP1^ interacts with the C-terminus of Epe1 through a region that is proximal to the JmjC domain. **A)** Western blots of *in vitro* binding assays to validate the interaction between Swi6 ^HP1^ and the Epe1 C-terminus. Increasing amounts of MBP-Epe1^434-948^ protein is added while maintaining a fixed amount of 3X-FLAG-Swi6^HP1^on beads. MBP-Epe1^434-948^ exhibits an increase in its interaction with Swi6 ^HP1^ compared to full-length Epe1. **B)** Increasing amounts of MBP-Epe1^434-600^ protein is added while maintaining a fixed amount of 3X-FLAG-Swi6^HP1^ on beads. MBP-Epe1^434-600^ exhibits an increase in binding to Swi6 ^HP1^ compared to full-length Epe1. **C)** A recombinant C-terminal fragment of Epe1, MBP-Epe1^434-948^ is incubated with fission yeast cell extracts expressing the N-terminal half of the protein, 3X FLAG-Epe1^1-434^ and 3X FLAG-Epe1^1-434^ H297A. Extracts were derived from *swi6+* cells. **D)** Same as C, extracts were derived from *swi6Δ* cells. **E)** Phenotype assays of epigenetic inheritance as a measure of Epe1 activity. Cells expressing Epe1^434-948^ were plated on –tetracycline and +tetracycline medium. Cells are initially red during establishment. However, despite the absence of a JmjC domain, cells partially turn white on +tetracycline medium. **F)** Quantification of red or sectored colonies comparing *epe1Δ* and Epe1^434-948^ expressing cells. The JmjCΔ mutant allele has 30% fewer sectored colonies compared to *epe1Δ* cells.

Although the Swi6^HP1^ binding site lies outside the confines of the JmjC domain of Epe1, point mutations within the putative catalytic perturb a direct interaction between the two proteins (**Figure 2A**). We speculated that there may exist an auto-inhibitory conformation of Epe1 where the N-terminus of the protein (1-434 amino acids) interacts with its C-terminal counterpart (434-948 amino acids) in *cis.* To test this model, we expressed the N-terminal half of Epe1 containing the JmjC domain fused to a 3X-FLAG epitope tag in fission yeast cells (3XFLAG-Epe1 1-434 amino acids). We expressed both the wild-type and the Fe (II) binding mutant allele, Epe1 H297A. We prepared cell extracts and directly incubated the lysate with a recombinant C-terminal Epe1 fragment, MBP-Epe1^434-948^ immobilized on an amylose resin. Compared to beads only controls where the amylose resin was directly incubated with cell extracts, we found that MBP-Epe1^434-948^ is able to pull-down the N-terminal half of Epe1 in *trans* (**Figure 3C)**. This interaction persists both in the case of the wild-type and JmjC mutant protein. Furthermore, we also observed this *trans* interaction in *swi6Δ* cells suggesting that the binding between the N and C terminal halves of Epe1 is not mediated by Swi6^HP1^ (**Figure 3D)**. These results support the existence of a *cis* interaction between the N- and C-terminus of Epe1 in the context of the full length protein raising potential mechanistic questions of how this auto-inhibitory conformation might be regulated.

One prediction that emerges from our biochemical analyses is that expressing the Epe1 C-terminus (Epe1^434-948^) alone might oppose heterochromatin assembly through a direct interaction with Swi6^HP1^. As previously described, we used a reporter gene assay where a TetR-Clr4-I fusion protein initiates heterochromatin establishment in an inducible manner. We expressed Epe1^434-948^ (JmjCΔ) in this reporter strain. Surprisingly, this mutant protein which completely lacks the JmjC domain can reverse epigenetic silencing during maintenance. This reversal leads to a greater proportion of cells that turn white upon +tetracycline addition (**Figure 3E**). The Epe1 JmjCΔ mutant produces a phenotype that is substantially different compared to Epe1 co-factor binding mutants or *epe1Δ* cells. We quantified the number of red or sectored colonies in the Epe1 JmjCΔ mutant compared to *epe1Δ* and wild-type cells. Cells that express a full-length wild-type copy of Epe1 turn white on +tetracycline medium and show no trace of red, pink or sectored colonies. We observed a three-fold reduction in the number of red or sectored colonies in the Epe1 JmjCΔ mutant compared to Epe1 null cells (**Figure 3F**). Therefore, the Epe1^434-948^ mutant is a hypomorphic allele which partially retains wild-type levels of Epe1 activity. We also observed clone to clone variability during maintenance with some cells resembling *epe1Δ* cells (data not shown). Differences in Epe1^434-948^ expression levels between strains did not seem to account for the variable phenotypes we observed in our silencing assays.

### H3K9 methylation stimulates complex formation between Epe1 and Swi6^HP1^

We speculated that a heterochromatin specific trigger might regulate the auto-inhibited conformation of Epe1 and potentially influence its interaction with Swi6^HP1^. We expressed Epe1 fused to a 3XFLAG epitope tag in cells which lack the H3K9 methyltransferase, Clr4^Suv39h^ or strains where the wild-type H3 allele is replaced with an H3K9R mutant. We performed a co-immunoprecipitation assay where we pull-down Epe1 with a FLAG antibody and measured its interaction with Swi6^HP1^. Although Epe1 interacts with Swi6^HP1^ in wild-type cells, this interaction is obliterated in both of the H3K9 methylation deficient mutant strains, *clr4Δ* and H3K9R mutants (**Figure 4A**). Furthermore, deleting histone deacetylases Sir2 or Clr3, both of which affect heterochromatin formation and Swi6^HP1^ localization, also resulted in a substantial decrease in the interaction between Epe1 and Swi6^HP1^ (**Figure 4B**). In contrast, deleting Mst2, a histone acetyltransferase that enhances heterochromatin formation, leads to no change in the interaction between wild-type Epe1 and Swi6^HP1^ and a loss of interaction in the Epe1 H297A mutant (**Supplementary Figure 4A**). Hence, our results indicate that H3K9 methylation and functional heterochromatin are pre-requisites for complex formation between Epe1 and Swi6^HP1^ in cells.

**Figure 4.**
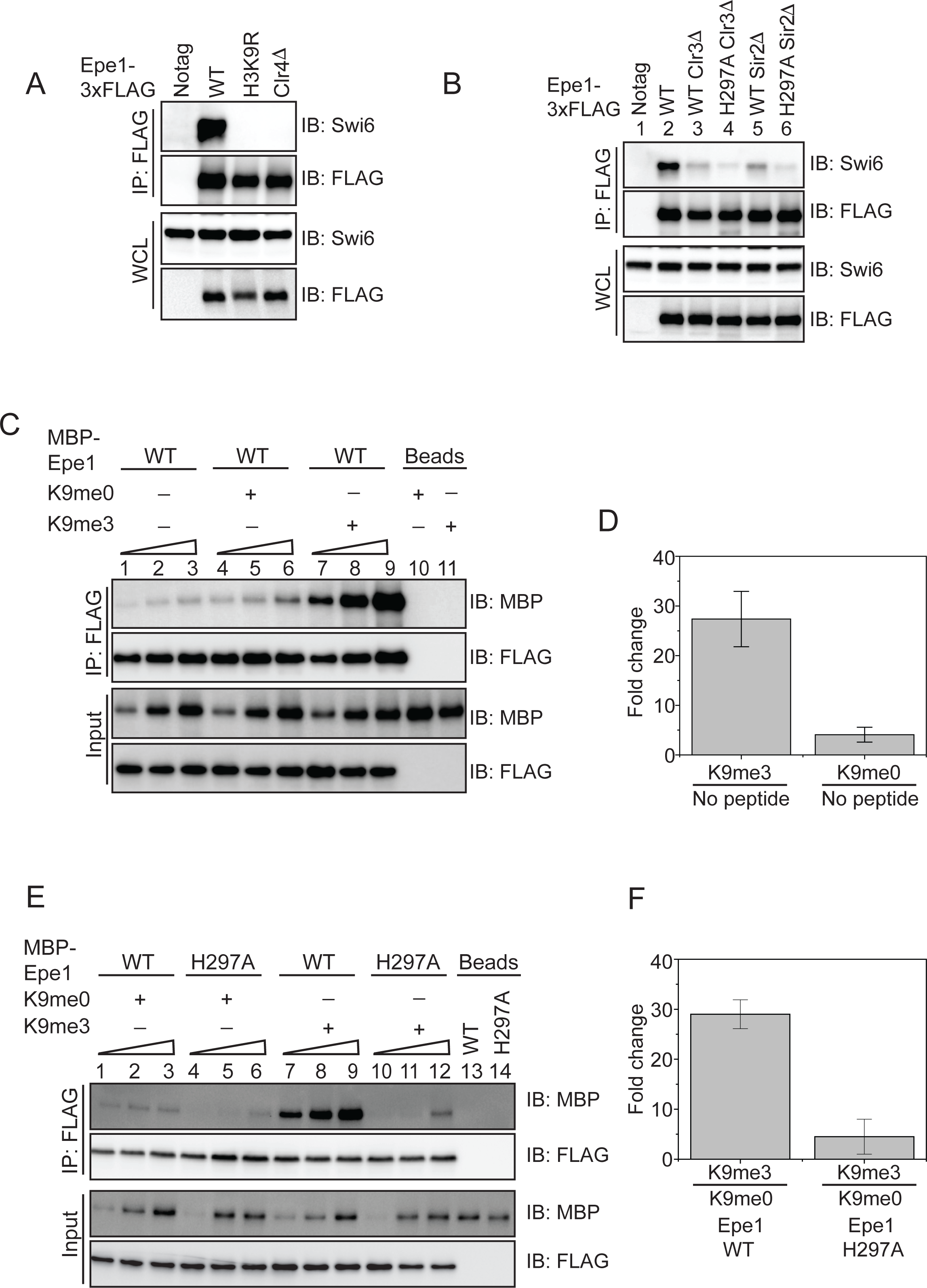
H3K9 methylation stimulates complex formation between Epe1 and Swi6^HP1^. **A)** Western blots of co-immunoprecipitation measurements reveal an interaction between Epe1-3XFLAG and Swi6^HP1^ in the case of wild-type cells which is completely absent in H3K9 methylation deficient cells (clr4Δ and H3K9R). **B)** Co-immunoprecipitation measurements reveal that the deletion of histone deacetylases Clr3 or Sir2 that disrupts heterochromatin formation also leads to a concomitant loss in the interaction between Epe1-3XFLAG and Swi6 ^HP1^. **C)** Western blots of *in vitro* binding assays between recombinant Epe1 and Swi6 ^HP1^ in the presence of histone H3 tail peptides (1-21 amino acids). Increasing amounts of wild-type MBP-Epe1 are added while maintaining a fixed amount of 3XFLAG-Swi6^HP1^ on beads. Experiments were performed in the presence of an unmethylated H3 peptide (H3K9me0) (lanes 4-6) or an H3K9 tri-methylated peptide (H3K9me3) (lanes 7-9). Epe1 exhibits a significant increase in its ability to interact with Swi6^HP1^ in the presence of an H3K9me3 peptide. **D)** Mean fold change in the interaction between Epe1 and Swi6^HP1^ in the presence of an H3K9me3 peptide versus an H3K9me0 peptide. Standard deviation represents the variation in the fold change values we measured across three different Epe1 concentrations in the *in vitro* binding assay. **E)** Western blots of *in vitro* binding assays between recombinant Epe1 WT and Epe1 H297A with Swi6^HP1^ in the presence of an H3K9me3 tail peptide. Increasing amounts of wild-type MBP-Epe1 and MBP-Epe1 H297A are added while maintaining a fixed amount of 3XFLAG-Swi6^HP1^ on beads in the presence of an H3K9me3 peptide or an unmodified H3K9me0 peptide. We observed a stimulation in the interaction between Epe1 WT and Swi6^HP1^ (lanes 7-9) but not in the case of Epe1 H297A and Swi6^HP1^ (lanes 10-12). **F)** Mean fold change in the interaction between Epe1 WT and Epe1 H297A with Swi6^HP1^ in the presence of an H3K9me3 peptide. Standard deviation represents the variation in the fold change values we measured across three different Epe1 concentrations in the *in vitro* binding assay.

We reconstituted this requirement for H3K9 methylation using binding assays as previously described. We supplemented our *in vitro* binding assays with an unmethylated histone H3 peptide (H3K9me0, H3 1-21 amino acids) or an H3K9 tri-methylated peptide (H3K9me3, H3 1-21 amino acids). Compared to reactions where no-peptide (lanes 1-3) or an unmethylated H3 peptide was added (lanes 4-6), we observed a substantial increase in the interaction between Epe1 and Swi6^HP1^ specifically in the presence of an H3K9me3 peptide (lanes 7-9) (**Figure 4C-D)**. An H3K9me2 peptide was also capable of stimulating the interaction between Epe1 and Swi6^HP1^ compared to reactions where no peptide was added or reactions where an unmodified peptide was used (**Supplementary Figure 4B-C)**. To test whether the stimulation in the interaction between Epe1 and Swi6^HP1^ is specific to H3K9 methylation, we carried out binding assays in the presence of an H3K4 tri-methylated peptide (H3K4me3). The addition of an H3K4me3 peptide fails to enhance complex formation between Epe1 and Swi6^HP1^ unlike the significant enhancement in binding we observed upon addition of an H3K9me3 peptide (**Supplementary Figure 4D-E**). Hence, the stimulatory effect we observed in our binding assays is specific to either H3K9me2 or H3K9me3.

Our earlier results reveal a severe reduction in the interaction between Epe1 H297A and Swi6^HP1^ compared to the wild-type protein (**Figure 2E**). We compared binding assays between MBP-Epe1 wild-type and MBP-Epe1 H297A with Swi6^HP1^ in the presence of an H3K9me3 peptide or an H3K9me0 peptide. Although we observed a strong stimulation in the interaction between wild-type Epe1 and Swi6^HP1^ (compare lanes 1-3 with 7-9), this stimulatory effect was significantly attenuated in the Epe1 H297A mutant (compare lanes 4-6 with 10-12) (**Figure 4E-F**). These results suggest that the JmjC mutant of Epe1 is preferentially maintained in an auto-inhibited conformation that is refractory to any interaction with Swi6^HP1^ in the presence or absence of H3K9 methylation.

One possibility is that Swi6 undergoes a conformational change that relieves auto-inhibition upon binding to an H3K9me3 peptide. To test this hypothesis, we purified a mutant Swi6^HP1^ protein with a mutation in its chromodomain (Canzio et al., 2011). A tryptophan to alanine mutation (W104A) within the Swi6^HP1^ chormodomain causes a significant reduction in H3K9 methylation binding. We tested whether Swi6^HP1^ W104A protein binding to Epe1 can also be enhanced in the presence of an H3K9me3 peptide. We added increasing amounts of MBP-Epe1 while maintaining a fixed amount of 3X-FLAG Swi6^HP1^ W104A on beads. Despite Swi6^HP1^ being unable to bind to the modified H3 tail peptide, we observed a stimulation in the interaction Epe1 and the Swi6^HP1^ (**Supplementary Figure 4F**). Therefore, we concluded that Epe1 binds to an H3K9me3 peptide and undergoes a conformational change that enhances its interaction with Swi6^HP1^. These results might explain the requirement for heterochromatin in cells to license the interaction between Epe1 and Swi6^HP1^ (**Figure 4A**).

### Epe1 inhibits Swi6 dependent heterochromatin assembly through a non-enzymatic process

We hypothesized that Epe1 might outcompete other heterochromatin associated proteins that localize to sites of heterochromatin formation via the Swi6^HP1^. Genetic studies reveal that Epe1 and a histone deacetylase Clr3 have opposing effects on nucleosome turnover within heterochromatin domains (Aygun et al., 2013). One possibility is that the interaction between Epe1 and Swi6^HP1^ excludes heterochromatin agonists, such as Clr3 from sites of heterochromatin formation. We used a co-immunoprecipitation assay to measure the extent of interaction between Clr3 and Swi6^HP1^. Clr3 was fused to a 3X V5 epitope tag and expressed in a wild-type Epe1 and an Epe1 H297A background. We used a V5 antibody to pull-down Clr3 following which we detected Swi6^HP1^ using a primary antibody. We measured a weak interaction between Swi6^HP1^ and Clr3 in a wild-type Epe1 background that substantially increases in an Epe1 H297A strain background (**Figure 5A**). This positive change in the interaction between Clr3 and Swi6^HP1^ occurs in the absence of any increase in Swi6^HP1^ occupancy at the pericentromeric repeats (**Figure 5B**). These results are consistent with a model where Epe1 H297A is predominantly in an auto-inhibited state and fails to interact with Swi6^HP1^ thus allowing other heterochromatin associated proteins to take its place.

**Figure 5:**
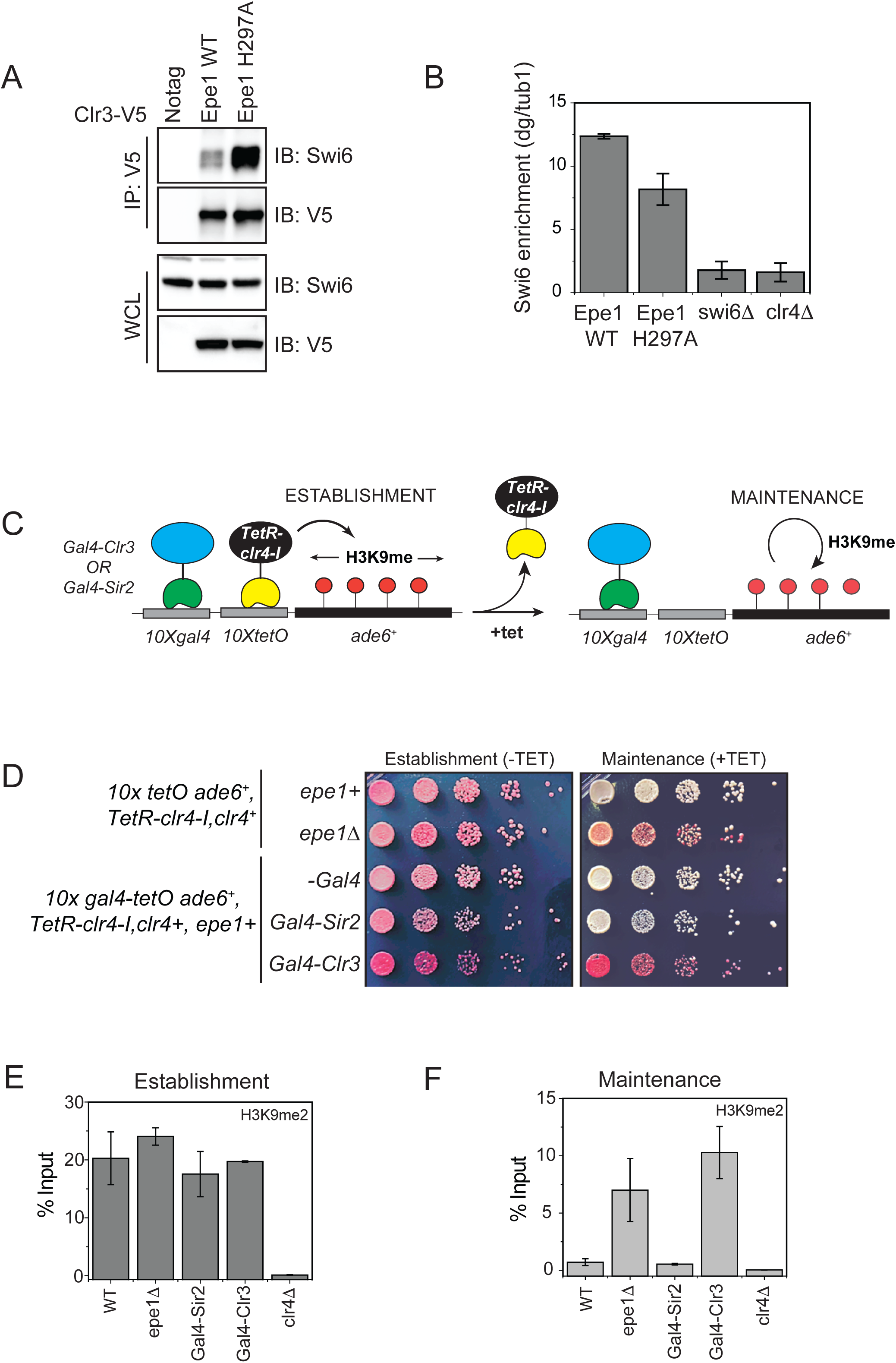
Tethering Clr3 to an ectopic site promotes epigenetic inheritance in the presence of Epe1. **A)** Epe1 displaces Clr3 from sites of heterochromatin formation. Western blots of a co-immunoprecipitation experiment reveals that Clr3-3X V5 interacts weakly with Swi6^HP1^ in a wild-type Epe1 background. This interaction increases significantly in an Epe1 H297A background. **B)** ChIP-qPCR measurements of Swi6^HP1^ occupancy at sites of constitutive heterochromatin in wild-type and Epe1 mutant cells. The increase in interaction between Swi6^HP1^ and Clr3 is independent of Swi6^HP1^ levels at sites of constitutive heterochromatin. **C)** A new reporter system to detect epigenetic inheritance in the presence of orthogonal chromatin effectors. Heterochromatin initiation depends on TetR-Clr4-I binding (-tetracycline). Orthogonal chromatin effectors can be recruited to the *10xgal4* DNA binding site via a Gal4 DNA binding domain (Gal4 DBD). The addition of tetracycline promotes TetR-Clr4-I dissociation to test for conditions where we can detect initiator independent maintenance of H3K9 methylation in the presence of an orthogonal chromatin effector. **D)** A color based assay to detect the establishment and maintenance of epigenetic states. The establishment of epigenetic silencing (-tetracycline) results in red colonies. Epigenetic inheritance, indicated by the appearance of red or sectored colonies, (+tetracycline) is altered in the presence of Gal4 fusions to HDAC proteins even in strains where Epe1 remains intact. **E)** ChIP-qPCR measurements of H3K9me2 levels at the ectopic site (*10X tetO ade6+*) during establishment (-tetracycline). (N=2). Error bars represent standard deviations. **F)** ChIP-qPCR measurements of H3K9me2 levels at the ectopic site (*10X tetO ade6+*) during maintenance (+tetracycline) (N=2). Error bars represent standard deviations.

We hypothesized that directly tethering Clr3 to sites of heterochromatin formation would nullify the non-enzymatic functions of Epe1. This would enable Clr3 to localize to sites of heterochromatin formation in a Swi6 independent manner. To test this model, we engineered synthetic heterochromatin domains where *10X Tet operator (10XTetO)* and *10X Gal4 DNA binding sites (10X Gal4)* are placed next to each other and inserted upstream of an *ade6+* reporter gene. This allows inducible heterochromatin formation via TetR-Clr4-I recruitment. The Gal4 DNA binding domain allows for the orthogonal recruitment of distinct chromatin effectors to the same ectopic site (**Figure 5C**). In the absence of any additional chromatin modifiers, TetR-Clr4-I recruitment to the newly engineered ectopic site (*10X Gal4-10X TetO-ade6+*) leads to the appearance of red colonies on –tetracycline containing medium. Upon addition of tetracycline, cells rapidly turn white consistent with a causal role for Epe1 in resetting epigenetic maintenance (**Figure 1C**). We fused the Gal4 DNA binding domain (Gal4 DBD) to two histone deacetylases, Clr3 (a class II histone deacetylase) and Sir2 (an NAD-dependent histone deacetylase) enabling these proteins to be constitutively tethered to the Gal4 DNA binding sequence. The TetR-Clr4-I fusion initiates heterochromatin formation and cells turn red on medium lacking tetracycline in the presence of Gal4-Clr3 or Gal4-Sir2 (**Figure 5D**). However, upon exposure to tetracycline, the constitutive tethering of Clr3 but not Sir2 promotes epigenetic maintenance. Cells exhibit a red and sectored phenotype in cells where Clr3 is artificially tethered, despite Epe1 still being fully active in these cells (**Figure 5D**). The phenotypes in cells where Gal4-Clr3 is tethered are remarkably similar to cells that lack Epe1 (*epe1Δ*). We measured changes in H3K9me2 levels associated with the *ade6+* reporter gene using ChIP-qPCR. Cells maintain high H3K9me2 levels in the presence of Gal4-Clr3 before and after tetracycline (**Figure 5E-F**). These results suggest that constitutively tethering Clr3 to sites of heterochromatin formation is sufficient to oppose Epe1 activity resulting in increased maintenance of H3K9 methylation after +tetracycline addition.

Tethering Gal4-Clr3 in the absence of TetR-Clr4-I causes no change in reporter gene silencing or H3K9me2 levels at the ectopic site (**Supplementary Figure 5A-C**). Hence, Gal4-Clr3 cannot initiate H3K9 methylation *de novo* in fission yeast (**Supplementary figure 5B-C)**. This lack of *de novo* silencing is consistent with the notion that HDAC proteins in fission yeast collaborate with H3K9 methyltransferases to establish epigenetic silencing. Furthermore, expressing Clr3 minus the Gal4 DBD fusion (Clr3 ΔGal4) leads to a loss of on +tetracycline containing medium. Therefore, Clr3 must be recruited and tethered in *cis* to repel the anti-silencing effects of Epe1 (**Supplementary Figure 5A)**. H3K9me2 levels are high during establishment but completely absent during maintenance (**Supplementary Figure 5B-C**). Hence, the sequence-specific recruitment of Clr3, rather than protein dosage, facilitates H3K9 methylation maintenance. Therefore, Clr3 recruitment functions in *cis* to maintain silent epigenetic states and antagonize Epe1 activity. This property of heterochromatin maintenance is RNAi independent since cells continue to exhibit a red or sectored appearance in a Dicer deficient background (*dcr1Δ*) (**Supplementary Figure 5D**).

The read-write activity of Clr4^Suv39h^ is essential for the inheritance of silent epigenetic states in a sequence-independent manner (Audergon et al., 2015; Ragunathan et al., 2015). This H3K9 methylation dependent positive feedback loop is disrupted in a Clr4^Suv39h^ chromodomain mutant (Zhang et al., 2008). To test whether the chromodomain is essential for maintenance when Clr3 is tethered, we replaced the wild-type allele of Clr4^Suv39^ with a Clr4 mutant that lacks the chromodomain (*clr4ΔCD)*. Cells that are initially red in –tetracycline medium turn white on +tetracycline medium in a *clr4ΔCD* expressing mutant (**Supplementary Figure 5E**). H3K9me2 levels in *clr4ΔCD* mutants are similar to that of wild-type cells during establishment. However, H3K9 methylation is absent upon +tetracycline addition in *clr4ΔCD* expressing strains. Hence, the inheritance of H3K9 methylation depends on the read-write activity of Clr4^Suv39h^ despite Clr3 being constitutively tethered (**Supplementary Figure 5F-G**).

## DISCUSSION

Auto-inhibition is a widely used strategy that proteins use to activate their latent functions in response to specific cellular or environmental signals (Pufall and Graves, 2002). The enzymatic and non-enzymatic properties of proteins can be regulated by internal control mechanisms that act in *cis.* For example, in the case of the chromatin remodeler ALC1, poly ADP ribosylation in response to DNA damage releases the macrodomain of the protein from an auto-inhibited state and activates its latent ATPase dependent nucleosome remodeling function (Lehmann et al., 2017; Singh et al., 2017). The non-catalytic heterochromatin associated protein Swi6^HP1^ is auto-inhibited by a histone H3 mimic sequence (Canzio et al., 2013). Release from auto-inhibition switches the protein to a spreading competent state. In our study, we demonstrate that a major antagonist of Swi6^HP1^ activity, a putative histone demethylase called Epe1, is also regulated by auto-inhibition. Remarkably, H3K9 methylation serves as a shared trigger between Epe1 and Swi6^HP1^ that relieves both proteins from auto-inhibition. In the absence of H3K9 methylation in cells, the two proteins fail to form a stable complex despite their ability to interact directly with each other.

We hypothesize that an interaction between the C-terminus of Epe1 and the putative catalytic JmjC domain preferentially maintains the protein in an auto-inhibited state. Alanine substitutions of nearly every Fe (II) or α−ketoglutarate binding residue obliterates the interaction between Epe1 and Swi6^HP1^. Release from auto-inhibition is H3K9 methylation dependent which confines complex formation between Epe1 and Swi6^HP1^ to a heterochromatin specific context (**Figure 6**). Swi6^HP1^ is dynamic and undergoes rapid exchange on the millisecond timescale between the free and H3K9 methylation bound state (Cheutin et al., 2004; Cheutin et al., 2003). One outcome of this H3K9 methylation dependent auto-inhibitory mechanism is that Epe1 selectively interacts with heterochromatin bound Swi6^HP1^ molecules as opposed to freely diffusing Swi6^HP1^ proteins. Given that Epe1 is expressed at levels that are at least 20-fold lower than Swi6 (data not shown), a most likely scenario is that a direct interaction would effectively titrate Epe1 away from sites of heterochromatin formation. By enforcing an H3K9 methylation dependent mode of interaction, Epe1 selectively targets a sub-population of Swi6^HP1^ molecules that are bound to sites of H3K9 methylation and have a causal role in heterochromatin formation.

**Figure 6:**
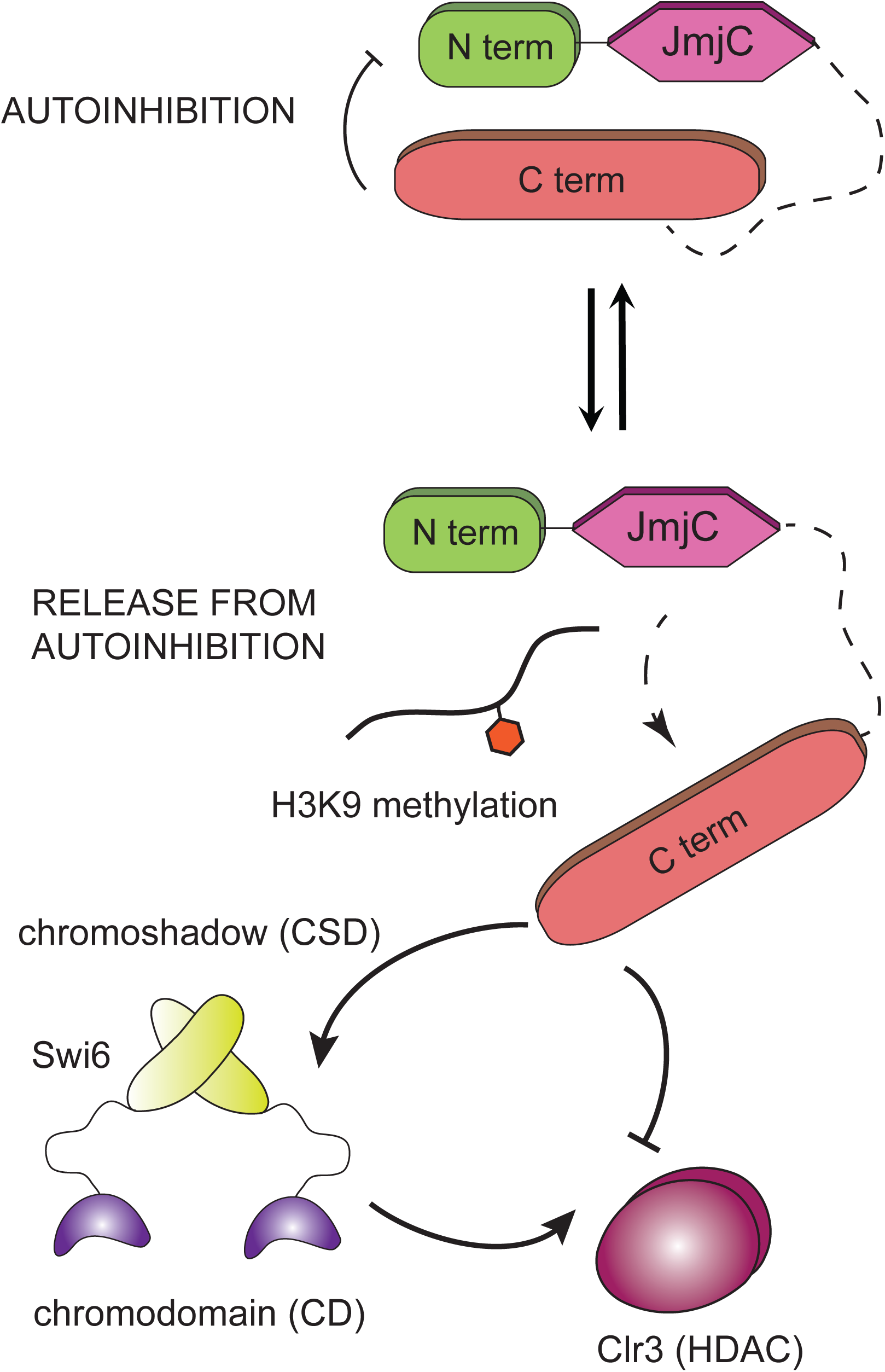
An auto-inhibited state of Epe1 regulates its non-enzymatic functions. Epe1 is in an auto-inhibited state that attenuates its interaction with Swi6^HP1^. Point mutations within the catalytic JmjC domain of Epe1 stabilize the auto-inhibited conformation which abolishes its binding to Swi6^HP1^. H3K9 methylation relieves auto-inhibition which has a stimulatory effect on the interaction between Epe1 and Swi6^HP1^. The interaction between Epe1 and Swi6^HP1^ displaces histone deacetylases such as Clr3 from sites of heterochromatin formation. Hence a competitive mode of binding which is triggered in the presence of H3K9 methylation defines the non-enzymatic functions of a putative histone demethylase.

We have identified the molecular basis for how Epe1 impairs epigenetic inheritance through a non-enzymatic mechanism. Our studies are consistent with earlier work demonstrating that Epe1 and the class II histone deacetylase, Clr3 have opposing functions that regulate heterochromatin assembly (Aygun et al., 2013; Shimada et al., 2009). Our findings reveal that these effects are not indirect but reflect a direct trade-off in protein-protein interactions involving Swi6^HP1^. Swi6^HP1^ occupancy at sites of heterochromatin formation is rate limiting with only a small fraction of the total protein being heterochromatin bound at any given time. Hence, the ability of Epe1 to interact with Swi6^HP1^ displaces other factors that promote heterochromatin assembly. Our studies show that the histone deacetylase Clr3 is exquisitely sensitive to the interaction between Epe1 and Swi6^HP1^ (**Figure 5A**). It is likely that this balance of protein-protein interactions is tunable via post-translational modifications. For example, Swi6^HP1^ phosphorylation compromises Epe1 binding and promotes histone deacetylase recruitment to sites of heterochromatin formation (Shimada et al., 2009). Our studies demonstrate that altering the balance of Swi6^HP1^ dependent protein-protein interactions profoundly affects the stability and heritability of silent epigenetic states.

Tethering Clr3 at an ectopic site renders heterochromatin refractory to the anti-silencing effects of Epe1. Our observations are in part, similar to previous findings relating to the fission yeast mating type locus where DNA binding proteins, Atf1 and Pcr1 recruit Clr3 to maintain epigenetic silencing following an RNAi dependent initiation mechanism (Wang and Moazed, 2017; Yamada et al., 2005). We also recapitulate a unique requirement for the sequence dependent recruitment of HDAC proteins in *cis* in order to facilitate the inheritance of silent epigenetic states despite the presence of anti-silencing factors such as Epe1. We favor a model where histone deacetylases such as Clr3 create a chromatin environment that promotes the read-write activity of Clr4^Suv39h^. Epe1 prevents Clr3 mediated histone hypoacetylation by forming an inhibitory complex with Swi6^HP1^ in a heterochromatin restricted genomic context. In addition Epe1 also directly interacts with members of the SAGA complex which promotes histone acetylation and transcription within heterochromatin repeats (Bao et al., 2019). Our results in conjunction with these studies collectively highlights the diverse non-enzymatic pathways that Epe1 might exploit to oppose Swi6^HP1^ mediated heterochromatin assembly.

The question of whether Epe1 harbors any histone demethylase activity remains unanswered. Inspired by earlier work on the *Drosophila* dKDM4A protein where HP1 binding stimulates demethylase activity, we attempted to reconstitute the enzymatic functon of Epe1 in the presence of Swi6^HP1^ {Lin, 2008 #1600}. However, our studies are unable to detect Epe1 mediated enzymatic demethylation in the presence or absence of Swi6^HP1^ (**Supplementary Figure 2F**). It is noteworthy that a histone demethylase like protein in *Neurospora*, DMM-1 shares many similarities with Epe1 and also surprisingly lacks any *in vitro* enzymatic activity (Honda et al., 2010). DMM-1 also interacts with the HP1 homolog in *Neurospora* leading us to speculate that this surprising mode of heterochromatin regulation we uncovered in the case of Epe1 might also extend to other fungal systems. Given that our Epe1 JmjCΔ allele only partially restores wild-type levels of Epe1 activity (**Figure 3E-F**), we speculate that there could potentially be unique substrates or conditions that could are required to *in vitro* reconstitute its enzymatic functions.

Our observations add to an expanding list of proteins that mimic histone modifiers but have built-in non-enzymatic functions that regulate the establishment and maintenance of epigenetic states. The *Drosophila* protein dKDM4A, is a prominent example of a histone demethylase with enzymatic and non-enzymatic functions (Colmenares et al., 2017). More extreme examples of histone modifying enzyme mimicry is seen in the case of JARID2, a subunit of the PRC2 complex that shares several characteristics with JmjC domain containing proteins but has a structural role in regulating PRC2 complex assembly (Kasinath et al., 2018; Son et al., 2013). Proteins involved in the *Arabidopsis* RNA dependent DNA methylation pathway, SUVH2 and SUVH9, resemble SET domain methyltransferases but lack enzymatic activity. Instead, both proteins recognize methylated DNA and are involved RNA pol V recruitment in plants to establish epigenetic silencing (Johnson et al., 2008; Law et al., 2013).

We hypothesize that Epe1 is a protein with dual-functionalities. Epe1 must interact with Swi6^HP1^ to be recruited to sites of heterochromatin formation. As a first approximation, the recruitment of Epe1 to sites of heterochromatin formation is alone sufficient to displace histone deacetylases and hasten the loss of transcriptional silencing. Hence, the non-enzymatic function of Epe1 serves as the first line of opposition to heterochromatin formation. This transient inhibition of heterochromatin assembly is likely to be followed by a slower enzymatic step where Epe1 may demethylate H3K9 methylated nucleosomes. This step would render heterochromatin assembly irreversible thus preventing its spreading and inheritance across generations. We envision that these distinct enzymatic and non-enzymatic properties of Epe1 oppose heterochromatin assembly on very different timescales. Our work highlights a unique mode of heterochromatin regulation where auto-inhibition regulates the non-enzymatic functions of putative histone demethylase and restricts its function to a heterochromatin specific context.

## ACKNOWLEDGEMENTS

We thank Ryan Baldridge, Patrick O’Brien and the Chromatin Club at the University of Michigan for insightful suggestions and critical feedback. We thank Hiten Madhani, Danesh Moazed and Songtao Jia for yeast strains. We also thank Danesh Moazed for sharing the Swi6 primary antibody which has been used for all of co-immunoprecipitation and chromatin immunoprecipitation (ChIP) experiments. We thank Nahid Iglesias for kindly sharing co-immunoprecipitation (co-IP) protocols prior to publication. This work was supported by startup funds from the Regents of the University of Michigan.

## MATERIALS AND METHODS

### Plasmids

Plasmids containing Epe1 wild-type and point mutants were constructed by modified existing pFA6a C-terminal tagging plasmids. Point mutations were introduced by designing primers using guidelines described in Quick Change mutagenesis protocols. A ligation independent cloning approach was used to construct pFastBac vectors containing wild-type Epe1 and Epe1 H297A mutant for recombinant protein expression and also for other MBP fusion constructs for *E.coli* expression. 3X FLAG Swi6^HP1^ and 3X FLAG Chp2^HP1^ was cloned into existing pGEX vectors downstream of the Prescission protease cleavage site using Gibson assembly. The construction of the *10X gal4-10X tetO-ade6^+^* plasmid involved modifying plasmids containing a 10X tetO sequence and sub-cloning Gal4 UAS sequences derived from a *Drosophila* pVALIUM 10X UAS vector. Vectors containing Gal4-Clr3 or Gal4-Sir2 were made using a modified pDual vector with an nmt1 promoter that enables facile incorporation of genetic payloads at the leu1 locus in fission yeast (Matsuyama et al., 2004). Further details regarding plasmid construction are readily available upon request.

### Strains

All strains were constructed using a PCR-based gene targeting approach (Bahler et al., 1998). In cases where we generated point mutations of *epe1*, we reintroduced the full length wild-type or mutant gene in *epe1Δ* strains. All strains were genotyped using colony PCR assays. We subsequently verified protein expression using western blots for each of the mutant strains. Strains with *10X gal4-10X tetO-ade6^+^* were constructed using an FOA selection strategy based on disrupting the endogenous *ura4* locus. Strains with Gal4-Clr3 or Gal4-Sir2 were made by digesting pDual vectors with a Not1 restriction enzyme followed by transformations and -LEU based selection. Other deletions of heterochromatin associated factors was achieved either by PCR-based gene targeting approaches or by a cross followed by random spore analysis and PCR based screening to select for colonies that harbored the reporter gene, All strains used in this study are listed in Table S1. Further details regarding strain construction are available upon request.

### Cell lysis, Co-immunoprecipitation and Western blotting

1.5 L of fission yeast cells cells were grown in YEA medium at 32 °C to an OD600 =3.5 and harvested by centrifugation. The cell pellets were washed with 10 ml TBS pH 7.5, re-suspended in 1.5 ml lysis buffer (30 mM Hepes pH 7.5, 100 mM NaCl, 0.25% Triton X-100, 5 mM MgCl2, 1 mM DTT) and cell suspension was snap-frozen into liquid nitrogen to form yeast “balls” and cryogenically ground using a SPEX 6875D Freezer/Mill. The frozen cell powder was thawed at room temperature and re-suspended in an additional 10 ml of lysis buffer with protease inhibitor cocktail and 1 mM PMSF. The cell lysates were subjected to two rounds of centrifugation at 18000 rpm for 5 and 30 mins in a JA-25.50 rotor (Beckman). Bradford assay was used to normalize protein levels for co-immunoprecipitation and immunoblot analysis. Protein G Magnetic Beads were pre-incubated with antibody for 4 h and cross-linked with 10 volumes of cross-linking buffer containing 20 mM DMP (3 mg DMP/ml of 0.2 M Boric Acid pH 9) for 30 min at room temperature by rotating. Cross linking was quenched by washing twice and incubated with 0.2 M ethanolamine pH 8 for 2 h at room temperature by rotating. The cell lysates were then incubated with antibody cross-linked beads for 3 hrs at 4 °C. Beads were washed 3 times in 1ml lysis buffer 5 mins each, then eluted with 500 ml of 10 mM ammonium hydroxide, the ammonium hydroxide was evaporated using speed vac (SPC-100H) for 5 h and re-suspended in SDS sample buffer. Samples were resolved on SDS-PAGE and transferred to PVDF membranes. Immunoblotting was performed by blocking PVDF membrane in Tris-buffered saline (TBS) pH 7.5 with 0.1% Tween-20 (TBST) containing 5% non-fat dry milk and subsequently probed with desired primary antibodies and secondary antibodies. Blots were developed by ECL method and detected with Bio-Rad ChemiDoc Imaging System.

### Chromatin Immunoprecipitation (ChIP)

Cells were grown till late log phase (OD 600-1.3 to 1.8) in yeast extract supplemented with adenine (YEA) or YEA containing tetracycline (2.5μg/ml) medium and fixed with 1% formaldehyde for 15 min at room temperature (RT). 130 mM glycine was then added to quench the reaction and incubated for 5 min at RT. The cells were harvested by centrifugation, and washed twice with TBS (50 mM Tris, pH 7.6, 500 mM NaCl). Cell pellets were resuspended in 300 μl lysis buffer (50 mM Hepes-KOH, pH 7.5, 100 mM NaCl, 1mM EDTA, 1% Triton X-100, 0.1% SDS, and protease inhibitors) to which 500 μL 0.5 mm glass beads were added and cell lysis was carried out by bead beating using Omni Bead Ruptor at 3000 rpm × 30sec × 10 cycles. Tubes were punctured and the flow-through was collected in a new tube by centrifugation which was subjected sonication to obtain the fragment sizes of roughly 100-500 bp long. After sonication the extract was centrifuged for 15 min at 13000 rpm at 4°C. The soluble chromatin was then transferred to a fresh tube and normalized for protein concentration by the Bradford assay. For each normalized sample, 25μL lysate was saved as input, to which 225 μL of 1xTE/1% SDS were added (TE: 50 mM Tris pH 8.0, 1 mM EDTA). Dynabeads Protein A were preincubated with Anti-H3K9me2 antibody (Abcam, ab1220). For each immunoprecipitation, 2 μg antibody coupled to 30μL beads was added to 400μL soluble chromatin, and the final volume of 500μL was achieved by adding lysis buffer. Samples were incubated for 2h at 4°C, the beads were collected on magnetic stands, and washed 3 times with 1 ml lysis buffer and once with 1 ml TE. For eluting bound chromatin, 100μL elution buffer I (50 mM Tris pH 8.0, 10mM EDTA, 1% SDS) was added and the samples were incubated at 65°C for 5 min. The eluate was collected and incubated with 150 μL 1xTE/0.67% SDS in the same way. Input and immunoprecipitated samples were finally incubated overnight at 65°C to reverse crosslink for more than 6 hours. 60μg glycogen, 100 μg proteinase K (Roche), 44 ul of 5M LiCl, and 250 ul of 1xTE was added to each sample and incubation was continued at 55°C for 1h. Phenol/chloroform extraction was carried out for all the samples followed by ethanol precipitation. Immuno-precipitated DNA was resuspended in 100μL of 10 mM Tris pH 7.5 and 50 mM NaCl and was used for qPCR (SYBR Green) using an Eppendorf Mastercycler Realplex. For extra crosslinking, prior to fixing with 1% formaldehyde, the cultures were incubated at 18°C for 2 hours in a shaking incubator. The cells were pelleted and resuspended in 4.5 ml of 1x PBS. To this 1.5mM EGS (Ethylene glycol bis[succinimidylsuccinate]), Pierce (Fisher) was added and the samples were incubated at RT for 20 min with mild shaking before adding 1 % formaldehyde. The samples were then processed as mentioned above.

### Recombinant protein purification from insect cells and *E.coli*

MBP-His-TEV-Epe1 was cloned into a pFastBac vector (Thermo Fisher Scientific) and used for Bacmid generation. Low-titer baculoviruses were produced by transfecting Bacmid into Sf21 cells using Cellfectin II reagent (Gibco). Full length *S. pombe* Epe1 protein (wild-type and mutant) were expressed in Hi5 cells infected by high titer baculovirus which were amplified from Sf21 cells. After 44 hrs of infection, Hi5 cells were harvested and lysed in buffer A (30 mM Tris-HCl (pH 8.0), 500 mM NaCl, 5 mM EDTA, 5 mM β-mercaptoethanol with protease inhibitor cocktails) using Emulsiflex-C3 (Avestin). The cleared cell lysate was applied to Amylose resin (New England Biolabs) followed by washing with buffer A and elution with buffer A containing 10 mM maltose. The N-terminal His-MBP tag can be removed by TEV protease cleavage which was used to evaluate protein solubility. Proteins were further purified using a Superdex 200 (GE Healthcare) size exclusion columns. The protein was concentrated up to in a storage buffer containing 30 mM Tris-HCl (pH 8.0), 500 mM NaCl, 30% Glycerol and 1 mM TCEP.

Proteins were expressed in BL21 (DE3) cells. Cells were grown to log phase at 37°C, cooled on ice, and induced with 0.3mM IPTG before incubation for 18h at 18°C. Pellets were suspended in Tris Buffered Saline (TBS) and frozen at −80°C until further use. For purification, cell pellets were thawed in lysis buffer (500mM NaCl, 50mM Tris pH 7.5, 10% glycerol) supplemented with protease inhibitor and cells were ruptured using sonicator. Cells debris was removed by centrifugation and the supernatant was incubated with proper beads for each protein for 3h at 4°C. We used a GST tag and glutathione beads (GST) beads for Swi6^HP1^ and Chp2^HP1^ purifications. We used an MBP tag and amylose resin for the purification of Epe1^434-948^ and Epe1^434-600^. After washing (lysis buffer was used as wash buffer), Epe1^434-948^ and Epe1^434-600^ was eluted with elution buffer (lysis buffer + 20mM maltose + 5mM EDTA). Swi6^HP1^ and Chp2^HP1^ were subject to overnight cleavage with Prescission protease.

### *In vitro* Binding Assay

*In vitro* binding assays were performed by immobilizing recombinant 3X FLAG-Swi6^HP1^ or 3X FLAG-Chp2^HP1^ on 25 μl of FLAG M2 beads which were incubated with three different concentrations of recombinant MBP fusion proteins in 600 μl binding buffer containing 20 mM Hepes pH 7.5, 150 mM NaCl, 5 mM MgCl2, 10% Glycerol, 0.25% Triton -X 100, 1 mM DTT. Reactions were incubated at 4 °C for 2 hrs and washed 3 times in 1ml washing buffer (20 mM Hepes pH 7.5, 150 mM NaCl, 5 mM MgCl2, 10% Glycerol, 0.25% Triton -X 100, 1 mM DTT) 5 min each, added 30μl SDS sample buffer followed by incubation at 95°C for 5 min. Proteins were separated through SDS-PAGE and transferred to PVDF membrane followed by incubation with anti-MBP monoclonal antibody (E8032S, NEB) and M2 Flag antibody (A8592, Sigma). Depending on the experiment, we added co-factors 100μM Ammonium iron (II) sulfate hexahydrate and 1 mM α-ketoglutarate or 5μg of H3 peptides (1-21 amino acids) with or without modifications. Western blot data for *in vitro* binding assays were analyzed using ImageJ software.

### Demethylase Assay

Mass spectrometry-based demethylase assays were performed using 5μg MBP-Epe1, 10μg Swi6, 20μM peptide (H3K9me3 peptide sequence used in this assay: NH2-ARTKQTAR(K9me3)STGGKA-amide), 50 mM HEPES (pH 7.5), 50 mM NaCl, 100 μM Ammonium iron (II) sulfate hexahydrate, 1 mM L-ascorbic acid, 1 mM α-ketoglutarate. Reaction mixtures were incubated at 37°C for 3 hrs, quenched with an equal volume of 1% trifluoroacetic acid, and stored at −20°C. In parallel, we also performed demethylase assays using equivalent amounts of purified JMJD2A (protein amounts equalized using SDS-PAGE gels). Samples were thawed and desalted using a ZipTip (Millipore). ZipTip was first equilibrated twice with wetting solution (50% Acetonitrile) and twice with equilibration solution (0.1% Trifluoroacetic acid) followed by 10μl samples bound to ZipTip and washed with washing solution (0.1% Trifluoroacetic acid) before elution with 4μl of 0.1% Trifluoroacetic acid/50% Acetonitrile. Matrix-assisted laser desorption ionization (MALDI) mass spectrometry was performed using a Waters Tofspec-2E in reflectron mode with delayed extraction (Department of Chemistry, University of Michigan).

### *In vitro* translation assays

To identify minimal Epe1 fragments that bind to Swi6^HP1^, Epe1 fragments were translated in *vitro* using TNT T7-coupled reticulocyte lysate (Promega) with ^35^S-labeled methionine (Roche). *In vitro* translated target proteins were incubated with Flag-tagged Swi6 at 4 °C for 20 min. Flag beads (Please find the product information) pre-equilibrated with buffer B containing 30 mM Tris-HCl (pH 8.0), 50 mM NaCl, 1 mM DTT and 0.1% NP-40 (w/v) were mixed and incubated at 4 °C for 45 min with rotation. The beads were washed three times with buffer B, and bead-bound proteins were separated by SDS-PAGE. Dried gels were analyzed by overnight exposure of a phosphor imager plate.

### *In vitro* binding assays using fission yeast cell extracts

100ml of fission yeast cells were grown in YEA medium at 32 °C to an OD600 =3 – 3.5 and harvested by centrifugation. The cell pellets were washed with 1 ml TBS pH 7.5 and resuspended in 600μL lysis buffer (30 mM Hepes pH 7.5, 100 mM NaCl, 0.25% Triton X-100, 5 mM MgCl2, 1 mM DTT) to which 500 μL 0.5 mm glass beads were added and cell lysis was carried out by bead beating using Omni Bead Ruptor at 3000 rpm × 30sec × 8 cycles. The cell extract was centrifuged for 20 min at 15000 rpm at 4°C and cell lysate incubated with beads pre-coupled with recombinant MBP-Epe1^434-948^ protein for 3h at 4°C, beads were washed 3 times with 1 ml lysis buffer and proteins were eluted by boiling the beads in SDS sample buffer. Proteins were resolved by SDS – polyacrylamide gel electrophoresis (SDS-PAGE) and analyzed by immunoblotting with appropriate antibodies.

---

**Supplemental Table S1.** Strains used in this study

| Strains | Strains description | Strain sources | Related to |
| --- | --- | --- | --- |
| KR18 | *h<sup>-</sup> leu1-32 ade6 M210 ura4Δ::10XTetO-ade6<sup>+</sup> clr4<sup>+</sup>, trp1:natMX6-clr4p-TetR-clr4ΔCD* | Moazed lab | Figure 1 A, 3E |
| KR33 | *h<sup>-</sup> leu1-32 ade6<sup>?</sup> ura4Δ::10XTetO- ade6<sup>+</sup> clr4<sup>+</sup>, trp1:natMX6-clr4p-TetR-clr4ΔCD, epe1Δ::kanMX6* | Moazed lab | Figure 1 A,C,D, 3E,F |
| KR283 | *h<sup>-</sup> leu1-32 ade6 M210 ura4Δ::10XTetO-ade6<sup>+</sup> clr4<sup>+</sup>, trp1:natMX6-clr4p-TetR-clr4ΔCD, 3XFLAG-epe1-hphMX6* | This study | Figure 1 A,C,D |
| KR303 | *h<sup>-</sup> leu1-32 ade6 M210 ura4Δ::10XTetO -ade6<sup>+</sup> clr4<sup>+</sup>, trp1:natMX6-clr4p-TetR-clr4ΔCD, 3XFLAG-epe1 H297A -hphMX6* | This study | Figure 1 A,C,D, 3E |
| KR844 | *h<sup>-</sup> leu1-32 ade6 M210 ura4Δ::10XTetO -ade6<sup>+</sup> clr4<sup>+</sup>, trp1:natMX6-clr4p-TetR-clr4ΔCD, 3XFLAG-epe1 Y307A -hphMX6* | This study | Figure 1 A,C,D, 3E |
| KR846 | *h<sup>-</sup> leu1-32 ade6 M210 ura4Δ::10XTetO -ade6<sup>+</sup> clr4<sup>+</sup>, trp1:natMX6-clr4p-TetR-clr4ΔCD, 3XFLAG-epe1 Y370A-hphMX6* | This study | Figure 1 A,C,D |
| KR24 | *h<sup>-</sup> leu1-32 ade6 M210 ura4Δ::10XTetO-ade6<sup>+</sup>, clr4Δ::kanMX6* | Moazed lab | Figure 1 C,D, 5E,F, S5B,C,F,G |
| KR817 | *h<sup>90</sup> ade6-M216 leu1-32 ura4-D18 mCherry-v5-Swi6 mNeonGreen-Epe1 WT-hphMX6* | This Study | Figure 1E |
| KR821 | *h<sup>90</sup> ade6-M216 leu1-32 ura4-D18 mCherry-v5-Swi6 mNeonGreen-Epe1 Y307A-hphMX6* | This Study | Figure 1 F |
| KR342 | *h<sup>+</sup> leu1-32 otr1R(SphI)::ura4<sup>+</sup> ura4-DS/E ade6 M210* | Moazed lab | Figure S1 B, 2A,B, S2A, 4A,B, S4A, 5A |
| KR471 | *h<sup>-</sup> leu1-32 epe1-3XFLAG-hphMX6* | This study | Figure S1 B, 2A,B, S2A, 4A,B, S4A, 5B |
| KR128 | *h<sup>-</sup> leu1-32 epe1-H297A-3XFLAG-hphMX6* | This study | Figure S1 B, 2A,B, S2A, 5B |
| KR769 | *h<sup>+</sup> leu1-32 ura4D18 imr1R(NcoI)::ura4<sup>+</sup> oriL ade6 M216 epe1 Y307A-3XFLAG-hphMX6* | This study | Figure S1 B, 2A,B, S2A |
| KR705 | *h<sup>+</sup> otr1R(SphI)::ura4<sup>+</sup> ura4-DS/E leu1-32 ade6 M210 epe1 Y370A -3XFLAG-hphMX6* | This study | Figure S1 B, 2A,B, S2A |
| KR132 | *h<sup>-</sup> leu1-32 epe1-3XFLAG-hphMX6 swi6Δ::kanMX6* | This study | Figure 2 B, S2A, 5B |
| KR947 | *h<sup>-</sup> leu1-32 3XFLAG-NLS-epe1 1-434 WT hphMX6* | This study | Figure 3 C |
| KR1210 | *h<sup>-</sup> leu1-32 3XFLAG-NLS-epe1 1-434 H297A hphMX6* | This study | Figure 3 C |
| KR1254 | *h<sup>-</sup> leu1-32 3XFLAG-NLS-epe1 1-434 WT hphMX6 swi6Δ::natMX6* | This study | Figure 3 D |
| KR1255 | *h<sup>-</sup> leu1-32 3XFLAG-NLS-epe1 1-434 H297A hphMX6 swi6Δ::natMX6* | This study | Figure 3 D |
| KR1197 | *h<sup>-</sup> leu1-32 ade6<sup>?</sup> ura4Δ::10XTetO- ade6<sup>+</sup> clr4<sup>+</sup>, trp1:natMX6-clr4p-TetR-clr4ΔCD, 3XFLAG-NLS-epe1 434-948 (JmjCΔ) #1* | This study | Figure 3 E, F |
| KR1198 | *h<sup>-</sup> leu1-32 ade6<sup>?</sup> ura4Δ::10XTetO- ade6<sup>+</sup> clr4<sup>+</sup>, trp1:natMX6-clr4p-TetR-clr4ΔCD, 3XFLAG-NLS-epe1 434-948 (JmjCΔ) #2* | This study | Figure 3 E, F |
| KR1199 | *h<sup>-</sup> leu1-32 ade6<sup>?</sup> ura4Δ::10XTetO- ade6<sup>+</sup> clr4<sup>+</sup>, trp1:natMX6-clr4p-TetR-clr4ΔCD, 3XFLAG-NLS-epe1 434-948 (JmjCΔ) #3* | This study | Figure 3 E, F |
| KR1227 | *h<sup>-</sup> leu1-32 3XFLAG-NLS-epe1 434-948 (JmjCΔ)* | This study | Figure S3C |
| KR228 | *h<sup>+</sup> otr1R(SphI)::ade6<sup>+</sup> ura4Δ18 leu1-32 ade6 M210 H3.1/H3.2/H3.3 K9R epe13X-FLAG-hphMX6* | Moazed lab | Figure 4A |
| KR130 | *h<sup>-</sup> leu-32 epe1 3X-FLAG-hphMX6 clr4Δ::kanMX6* | This study | Figure 2 B, S2A, 4A, 5B |
| KR724 | *h<sup>-</sup> leu-32 epe1 3X-FLAG hphMX6 clr3Δ::kanMX6* | This study | Figure 4 B |
| KR726 | *h<sup>-</sup> leu-32 epe1 H297A 3X-FLAG hphMX6 clr3Δ::kanMX6* | This study | Figure 4 B |
| KR728 | *h<sup>-</sup> leu-32 epe1 3X-FLAG-hphMX6 sir2Δ::kanMX6* | This study | Figure 4 B |
| KR730 | *h<sup>-</sup> leu-32 epe1 H297A 3X-FLAG-hphMX6 sir2Δ::kanMX6* | This study | Figure 4 B |
| KR732 | *h<sup>-</sup> leu1-32 ade6<sup>?</sup> ura4D18 mst2Δ::kanMX6 epe1 3X-FLAG-hphMX6* | This study | Figure S4 A |
| KR734 | *h<sup>-</sup> leu1-32 ade6<sup>?</sup> ura4D18 mst2Δ::kanMX6 epe1 H297A 3X-FLAG-hphMX6* | This study | Figure S4 A |
| KR906 | *h<sup>-</sup> leu1-32 epe1 3X-FLAG hphMX6 clr3 3X-V5-natMX6* | This study | Figure 5 A |
| KR908 | *h<sup>-</sup> leu1-32 epe1 H297A-3XFLAG-hphMX6 clr3 3X-V5-natMX6* | This study | Figure 5 A |
| KR675 | *h<sup>-</sup> ade6 M210 ura4Δ::10XGal4-10XTetO-ade6<sup>+</sup> natMX6-TetR-clr4-I, epe1<sup>+</sup>, leu1<sup>+</sup>:nmt1-gal4-sir2* | This study | Figure 5 D,E,F |
| KR677 | *h<sup>-</sup> ade6 M210 ura4Δ::10XGal4-10XTetO-ade6<sup>+</sup> natMX6-TetR-clr4-I epe1<sup>+</sup>, leu1<sup>+</sup>:nmt1-gal4-clr3* | This study | Figure 5 D,E,F, S5A,B,C,F,G |
| KR606 | *h<sup>-</sup> leu1-32 ade6 M210 ura4Δ::10XGal4-10XTetO-ade6<sup>+</sup> trp1:natMX6-TetR-clr4-I* | This study | Figure 5 D,E,F, S5B,C,F,G |
| KR643 | *h<sup>-</sup> leu1-32 ade6 M210 10XTetO- ade6<sup>+</sup> natMX6-TetR-Clr4-I, epe1Δ::kanMX6* | This study | Figure 5 D,E,F |
| KR1075 | *h<sup>-</sup> ade6 M210 ura4Δ::10XGal4-10XTetO-ade6<sup>+</sup> leu1<sup>+</sup>:nmt1-gal4-clr3* | This study | Figure 5 SA,B,C |
| KR1071 | *h<sup>-</sup> ade6 M210 ura4Δ::10XGal-10XTetO-ade6<sup>+</sup> natMX6-TetR-clr4-I epe1<sup>+</sup> leu1<sup>+</sup>:nmt1-clr3 (-gal4)* | This study | Figure 5D, S5 A |
| KR1067 | *h<sup>-</sup> ade6 M210 ura4Δ::10XGal-10XTetO-ade6<sup>+</sup> natMX6-TetR-clr4-I epe1<sup>+</sup> leu1<sup>+</sup>:nmt1-Gal4-clr3 hphMX6-NLS-clr4ΔCD #1* | This study | Figure S5 E,F,G |
| KR1068 | *h<sup>-</sup> ade6 M210 ura4Δ::10XGal-10XTetO-ade6<sup>+</sup> natMX6-TetR-clr4-I epe1<sup>+</sup> leu1<sup>+</sup>:nmt1-Gal4-clr3 hphMX6-NLS-clr4ΔCD #2* | This study | Figure S5 E |
| KR1103 | *h<sup>-</sup> ade6 M210 ura4Δ::10XGal4-10XTetO-ade6<sup>+</sup> natMX6-TetR-clr4-I epe1<sup>+</sup> leu1<sup>+</sup>:nmt1-gal4-clr3 dcr1Δ::kanMX6* | This study | Figure S5 D |

**Supplementary Figure 1:**
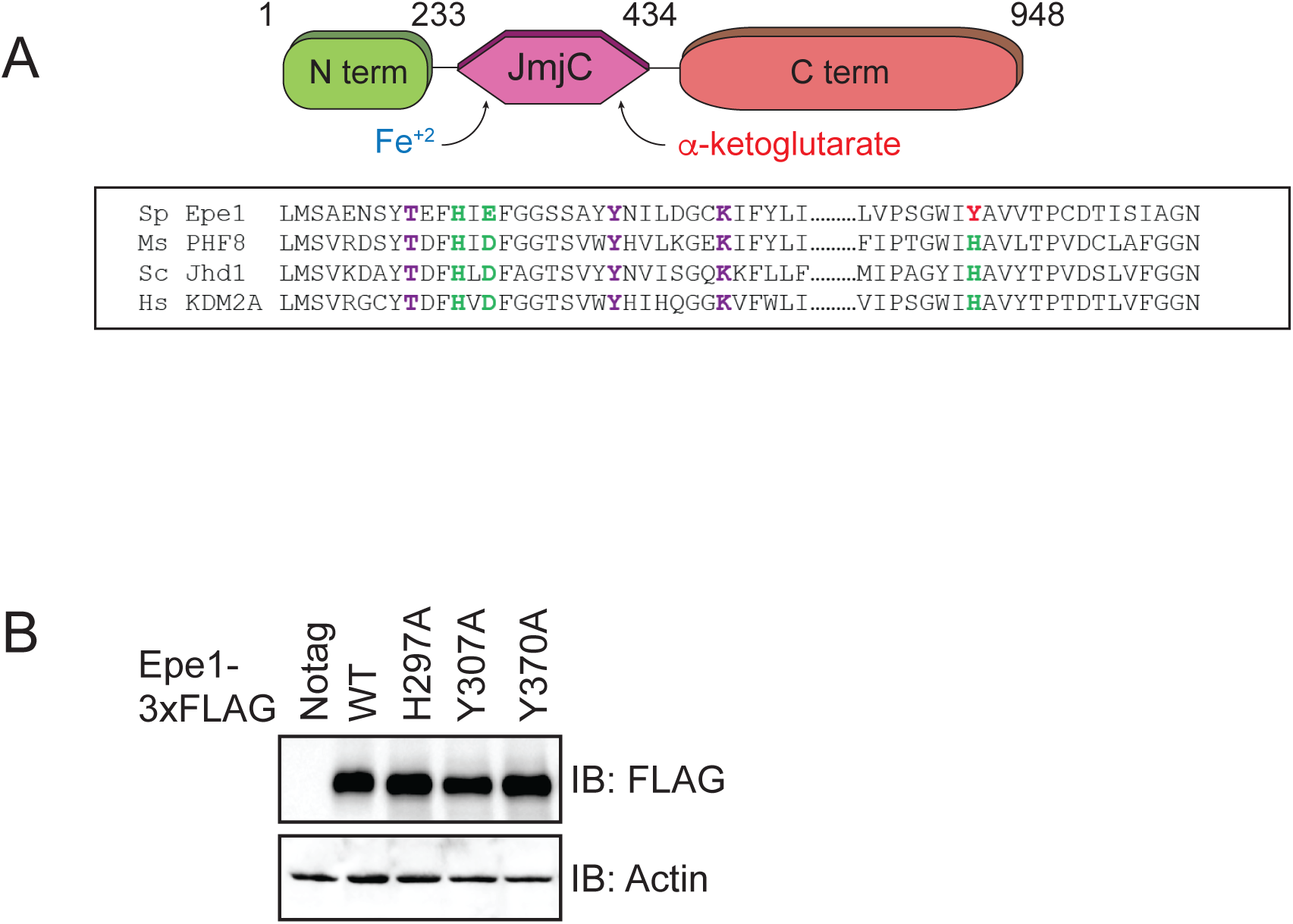
**A)** Sequence alignment of related JmjC domain containing proteins reveals the identity of conserved residues in Epe1 that are involved in iron (Fe^2+^) and α-ketoglutarate binding. **B)** Western blots comparing expression levels of Epe1 proteins. Wild-type Epe1-3X FLAG and Epe1 mutants exhibit equal levels of expression from the endogenous promoter. Actin levels are shown as a loading control.

**Supplementary Figure 2:**
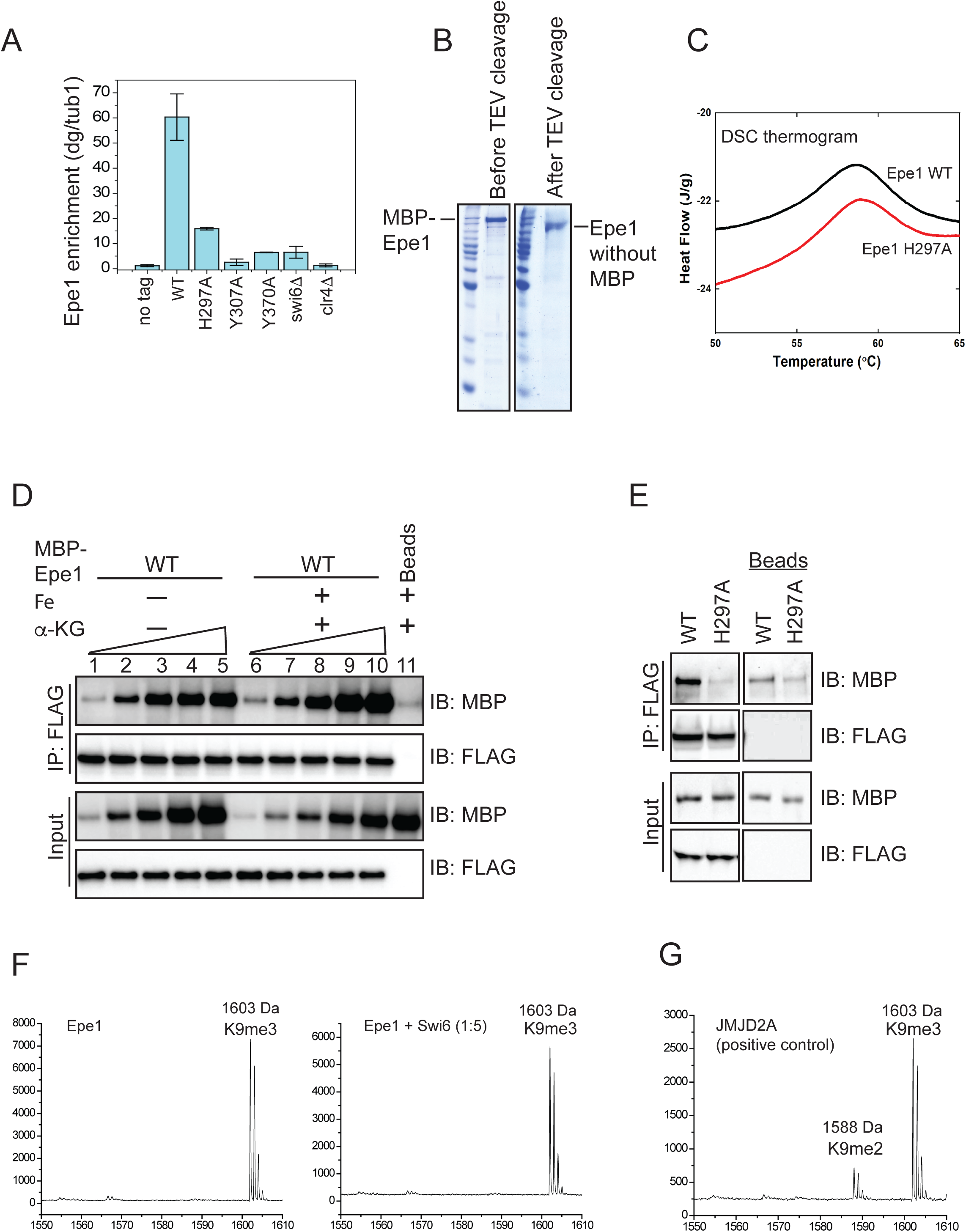
**A)** ChIP-qPCR measurements of Epe1 occupancy at the pericentromeric *dg* repeats. The use of additional cross-linkers improves crosslinking efficiency and traps transient interactions. The wild-type Epe1 protein is enriched at sites of heterochromatin formation whereas Epe1 JmjC mutants do not exhibit any significant enrichment. **B)** Recombinant Epe1 protein purification from insect cells before and after cleavage of the MBP tag. **C)** Differential scanning calorimetry (DSC) assays demonstrate that wild-type Epe1 and Epe1 H297A proteins exhibit similar denaturation temperatures implying that the proteins are folded and equally stable *in vitro*. The difference in peak intensities reflects slightly different protein amounts in the DSC **D)** *In vitro* binding assays between Epe1 and Swi6 in the presence of Fe (II) and α-ketoglutarate as co-factors. The presence of Fe (II) and α-ketoglutarate did not alter the binding between Epe1 and Swi6^HP1^ compared to reactions performed in the absence of any co-factors. **E)** *In vitro* binding experiments using recombinant wild-type and Epe1 H297A proteins with Swi6^HP1^ purified from fission yeast cells. 3X FLAG-Swi6^HP1^ was expressed at endogenous levels in fission yeast and purified under high salt conditions. The fission yeast Swi6^HP1^ protein exhibits the same binding preference for wild-type Epe1 as *E.coli* Swi6^HP1^. **F)** Mass spectrometry analysis of modified H3 peptides following an *in vitro* demethylase assay. Demethylase assays were carried out in the presence and absence of Swi6^HP1^. **G)** The positive control active demethylase, JMJD2A produces a 15kDa mass shift corresponding to the removal of a single methyl group.

**Supplementary Figure 3.**
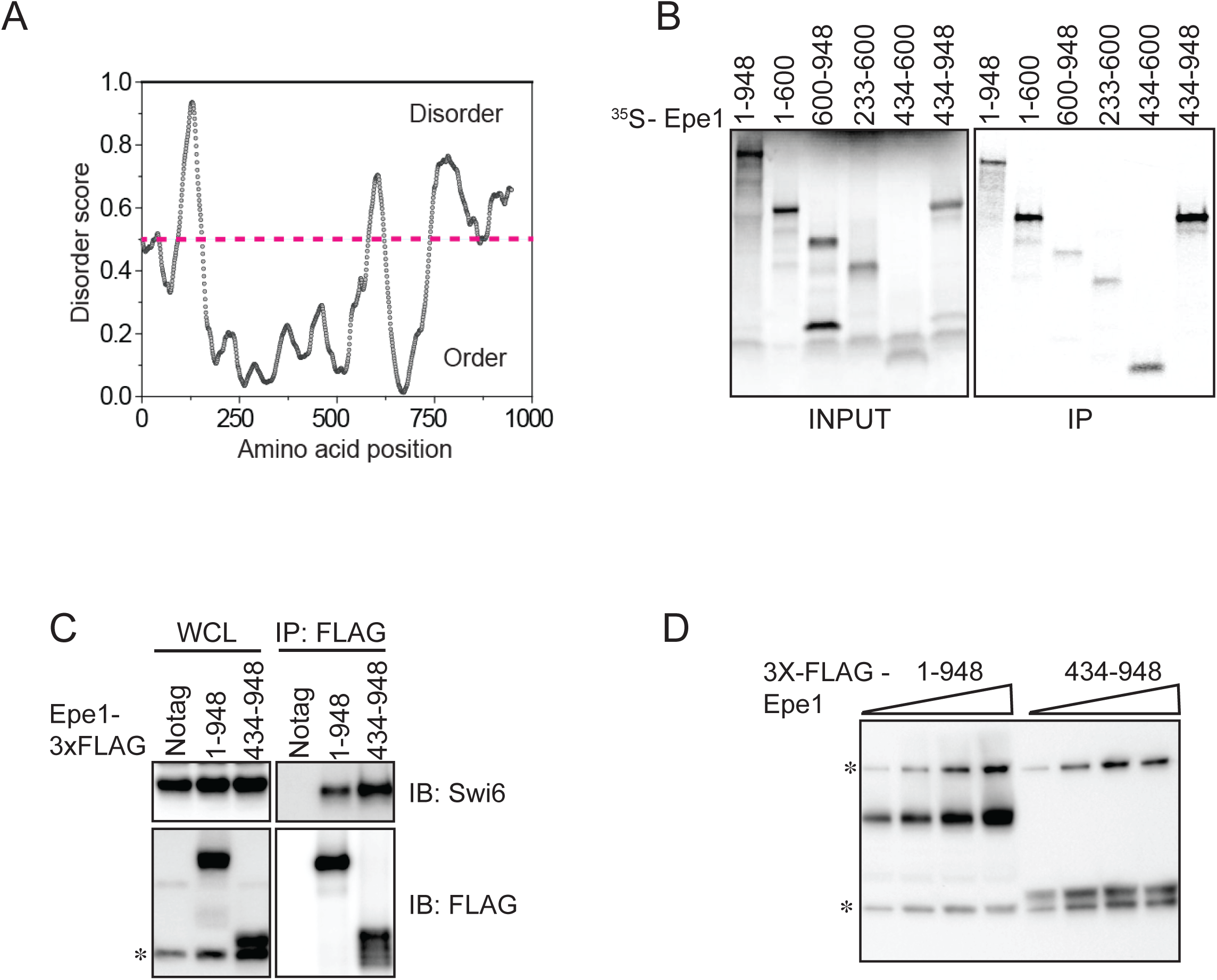
**A)** Using a computational disorder prediction program (http://www.pondr.com, VSL2) and a user specified cut-off, we defined ordered and disordered regions within Epe1. The JmjC domain spans 233-434 amino acids and emerges as one of two ordered domains in Epe1. The second ordered region within the C-terminus of Epe1 has no structural similarity to existing proteins. **B)** An *in vitro* translation assay using rabbit reticulocyte lysates to generate S^35^ labeled fragments of Epe1. Swi6^HP1^ was immobilized on beads and incubated with IVT extracts. Two Epe1 regions ranging from 1-600 amino acids and 434-948 amino acids emerge as the primary interacting fragments. **C)** Western blots of co-immunoprecipitation experiments to detect an interaction between Epe1^434-948^ and Swi6^HP1^.**D)** Western blots of full length Epe1 and Epe1^434-948^ fused to a FLAG epitope tag. The Epe1^434-948^ protein is expressed at levels that are 4-5 fold lower than full-length Epe1 which in part might explain its limited efficacy in cells. The asterisk in the figure denotes the non-specific FLAG antibody band.

**Supplementary Figure 4:**
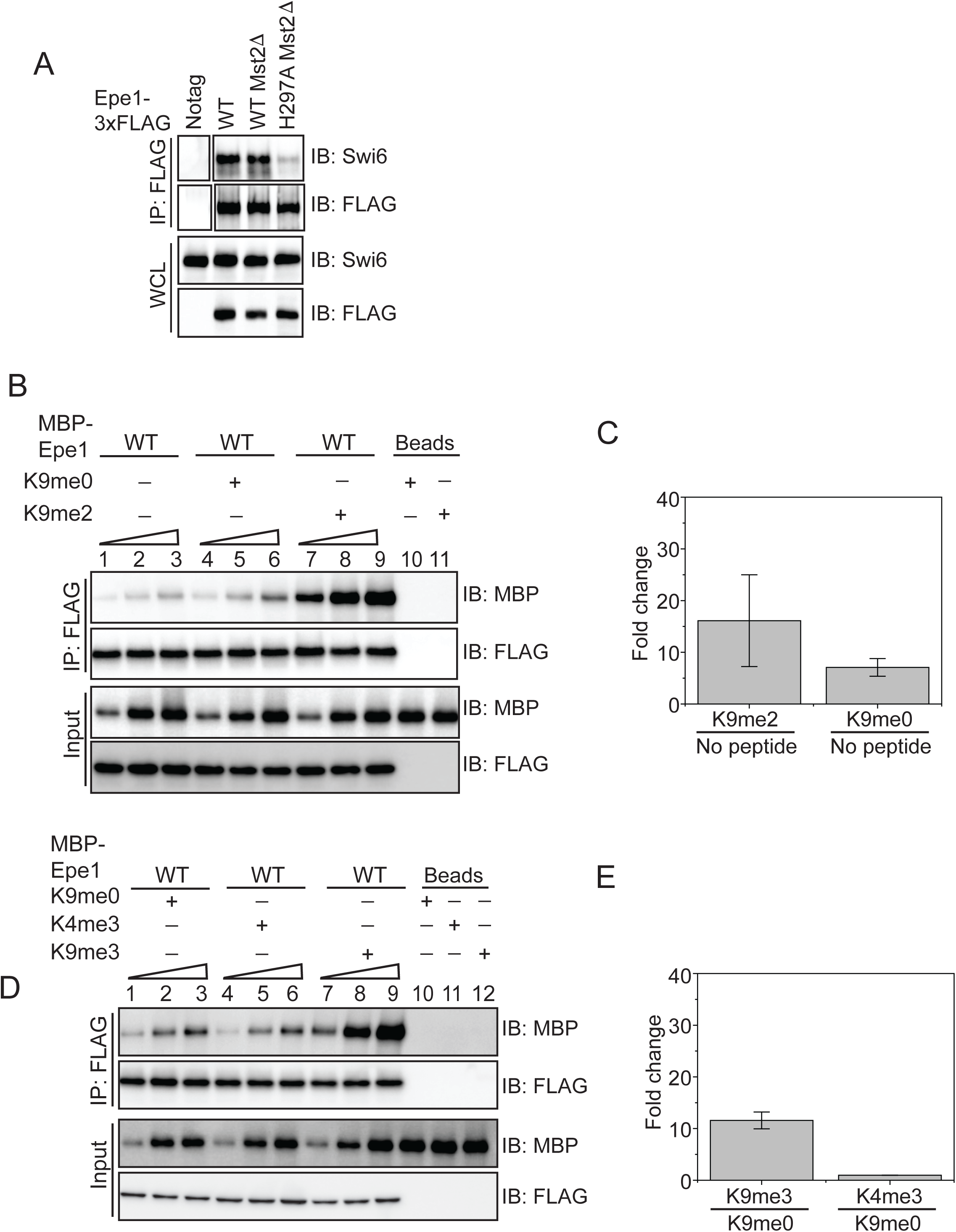
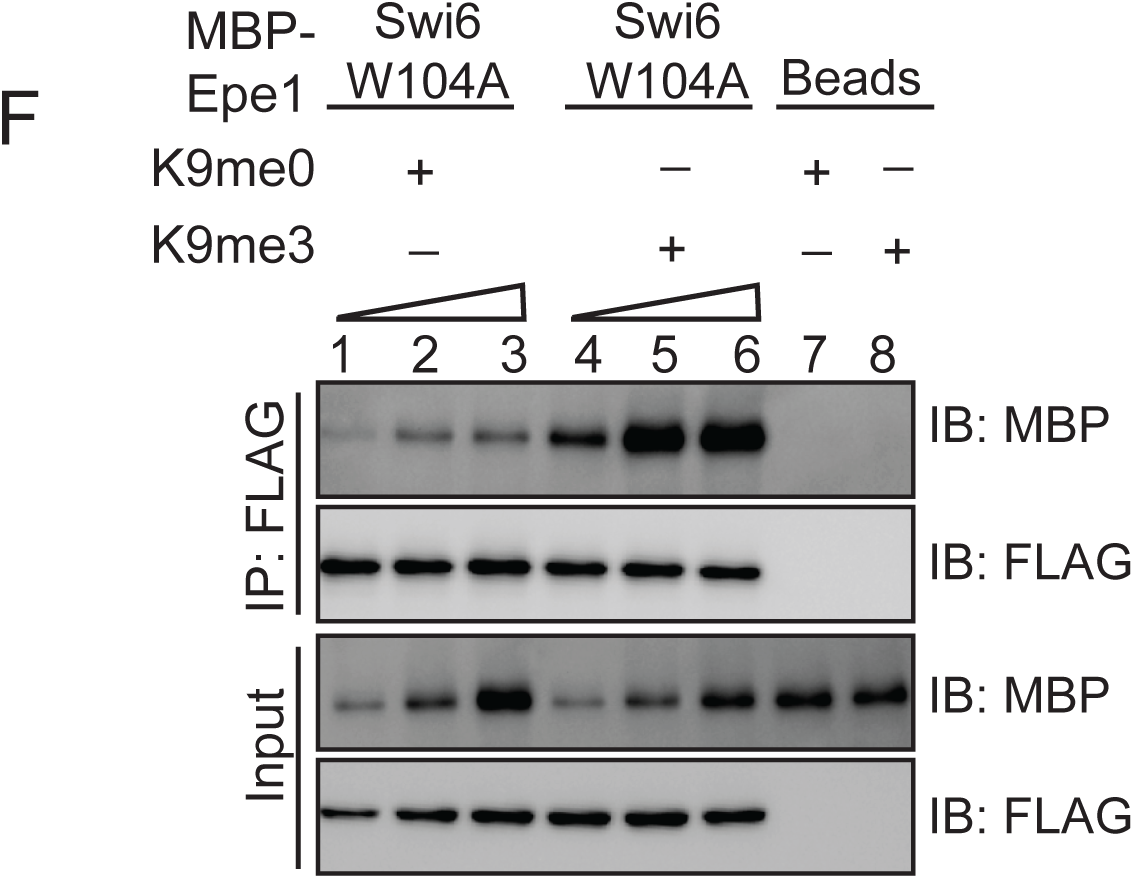
**A)** Deleting Mst2 does not alter the interaction between Epe1-3X FLAG and Swi6^HP1^. Western blots of a co-immunoprecipitation assay between Epe1 and Swi6 in an *mst2Δ* background. **B)** Western blots of *in vitro* binding assays between recombinant Epe1 and Swi6^HP1^ in the presence of histone H3 tail peptides (1-21 amino acids). Increasing amounts of wild-type MBP-Epe1 are added while maintaining a fixed amount of 3XFLAG-Swi6^HP1^ on beads. Experiments were performed in the presence of an unmethylated H3 peptide (H3K9me0) or an H3K9 di-methylated peptide (H3K9me2). Epe1 exhibits an increase in its ability to interact with Swi6^HP1^ in the presence of an H3K9me2 peptide similar to the type of stimulation we observed upon addition of an H3K9me3 peptide. **C)** Mean fold change in the interaction between Epe1 and Swi6^HP1^ in the presence of an H3K9me2 peptide versus an H3K9me0 peptide. Standard deviation represents the variation in the fold change values we measured across three different Epe1 concentrations in the *in vitro* binding assay. **D)** Western blots of *in vitro* binding assays between recombinant Epe1 and Swi6^HP1^ in the presence of differentially methylated histone H3 tail peptides (1-21 amino acids). Increasing amounts of wild-type MBP-Epe1 are added while maintaining a fixed amount of 3XFLAG-Swi6^HP1^ on beads. Experiments were performed in the presence of an unmethylated H3 peptide (H3K9me0), an H3K9 tri-methylated peptide (H3K9me3) or an H3K4 tri-methylated peptide (H3K4me3). Epe1 exhibits a significant increase in its ability to interact with Swi6^HP1^ in the presence of an H3K9me3 but not at H3K4me3 peptide. **E)** Mean fold change in the interaction between Epe1 and Swi6^HP1^ in the presence of an H3K9me3 peptide and an H3K4me3 peptide. Standard deviation reveals the variation in the fold change values we measured across three different Epe1 concentrations we used in our *in vitro* binding assay. **F)** Western blots of *in vitro* binding assays between recombinant Epe1 and Swi6^HP1^ W104A in the presence of histone H3 tail peptides (1-21 amino acids). Increasing amounts of wild-type MBP-Epe1 are added while maintaining a fixed amount of 3XFLAG-Swi6^HP1^ W104A on beads. Experiments were performed in the presence of an unmethylated H3 peptide (H3K9me0) or an H3K9 tri-methylated peptide (H3K9me3). Epe1 exhibits an increase in its ability to interact with Swi6^HP1^ W104A in the presence of an H3K9me3 peptide.

**Supplementary Figure 5:**
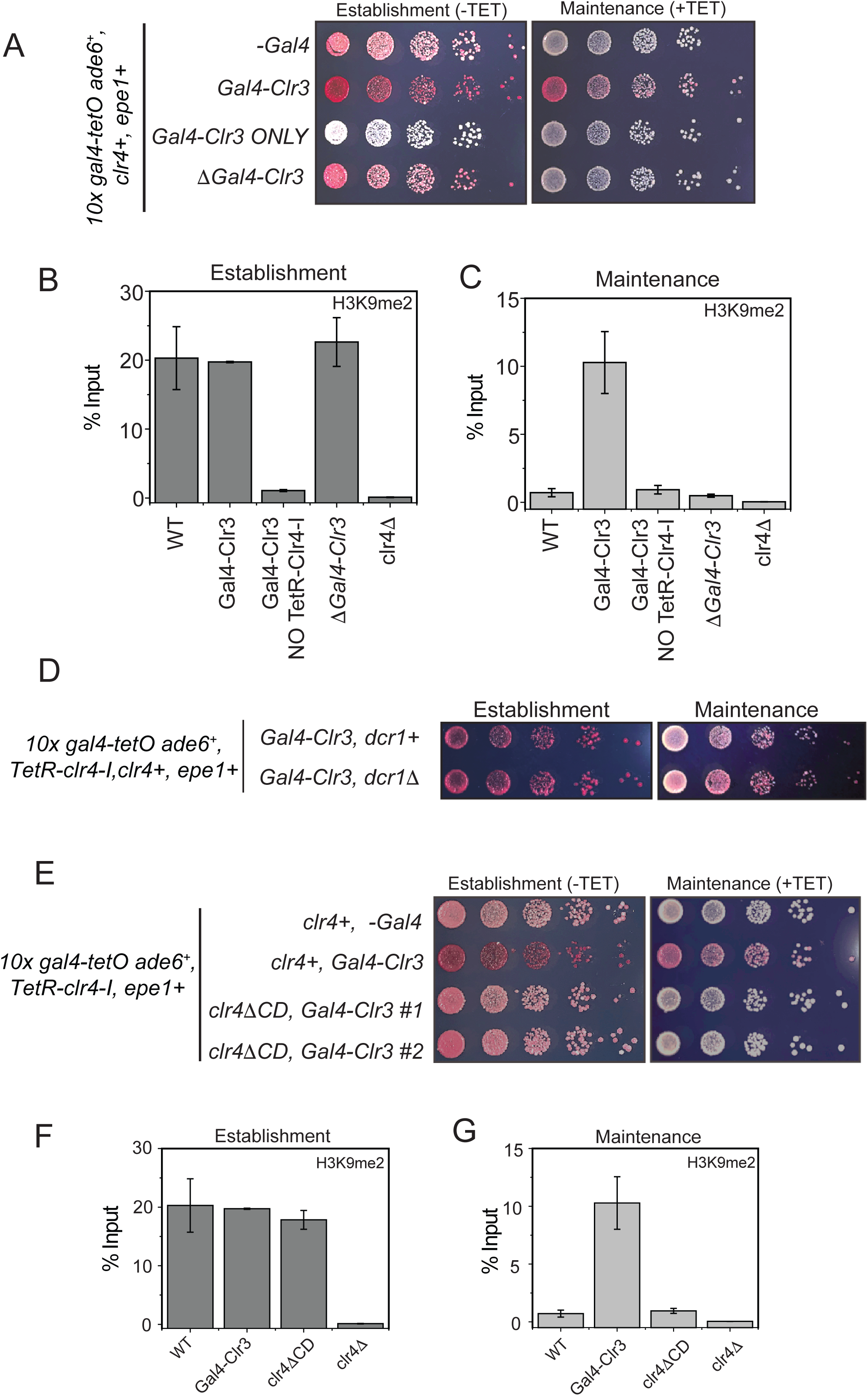
**A)** Color based assays provide a readout of the establishment and maintenance properties of a modified epigenetic inheritance reporter strain in the presence of Gal4-Clr3 fusion variants. Gal4-Clr3 fusions cannot initiate heterochromatin *de novo* in the absence of TetR-Clr4-I. Furthermore, Clr3 when overexpressed without the Gal4 DNA binding domain is unable to promote epigenetic inheritance (+tetracycline). **B)** ChIP-qPCR measurements of H3K9me2 levels at the ectopic site (*10X tetO ade6+*) during establishment (-tetracycline) in the presence of Gal4-Clr3 variants. ChIP-qPCR values from Figure 4C are plotted as a reference. **C)** ChIP-qPCR measurements of H3K9me2 levels at the ectopic site (*10X tetO ade6+*) during maintenance (+tetracycline) in the presence of Gal4-Clr3 variants (N=2). Error bars represent standard deviations. ChIP-qPCR values from Figure 4D are plotted as a reference. **D)** Color based assays provide a readout of the establishment and maintenance properties of a modified epigenetic inheritance reporter strain. *dcr1Δ* cells exhibit maintenance when Gal4-Clr3 is constitutively tethered at an ectopic site. Maintenance via Gal4-Clr3 recruitment is independent of the RNAi pathway.**E)** The deletion of the Clr4 chromdomain (*clr4ΔCD)* affects its read-write. Cells that are initially red during heterochromatin establishment (-tetracycline) turn white during maintenance (+tetracycline) in a *clr4ΔCD* background despite Clr3 being constitutively tethered. **F)** ChIP-qPCR measurements of H3K9me2 levels at the ectopic site (*10X tetO ade6+*) during establishment (-tetracycline) in cells that have a Clr4 chromdomain mutant (N=2). Error bars represent standard deviations. ChIP-qPCR values from Figure 5C are plotted as a reference. **G)** ChIP-qPCR measurements of H3K9me2 levels at the ectopic site (*10X tetO ade6+*) during maintenance (+tetracycline) in cells that have a Clr4 chromdomain mutant (N=2). Error bars represent standard deviations. Error bars represent standard deviations. ChIP-qPCR values from Figure 5D are plotted as a reference.

